# Illuminating spatial dynamics of glutamine metabolism with a sensitive genetically encoded biosensor

**DOI:** 10.64898/2026.01.15.699810

**Authors:** Jack M. Scully, Michael J. Sun, Siddharth Das, Nataly J. Arias, Edmund D. Kapelczak, Tiffany D. Robinson, Katie G. Vineall, Teagan S. Dean, Tara TeSlaa, Ajit S. Divakaruni, Danielle L. Schmitt

## Abstract

Glutamine is the most abundant amino acid in serum, used as a key nutrient by cells for protein synthesis, energy production, carbon and nitrogen metabolism, and cellular redox balance. The use of glutamine in the cell is highly compartmentalized, but the dynamics of glutamine metabolism across organelles and individual cells are not fully understood. To illuminate subcellular glutamine dynamics, we developed a green fluorescent protein-based intracellular glutamine optical reporter, iGlo. We find iGlo is sensitive and specific for glutamine and can be used to measure glutamine uptake, production, and consumption with high spatiotemporal resolution in multiple cell types. Furthermore, multiplexed imaging of iGlo with a lactate biosensor in single cells reveals the temporal crosstalk between glucose and glutamine metabolism to maintain energy homeostasis. Thus, iGlo enables the sensitive and precise study of compartmentalized glutamine dynamics and represents a new and enhanced tool for studying the spatiotemporal dynamics and regulation of metabolism.

## Introduction

Glutamine is the most abundant amino acid in serum, playing versatile roles in processes beyond its use as a proteinogenic amino acid including energy production, redox and pH homeostasis, and nucleotide, lipid, and amino acid biosynthesis^1^. Dysregulation of glutamine metabolism is also linked to pathophysiological states, including cancer, several types of which are reported to be dependent on glutamine to fuel metabolism and cell proliferation^2^. Glutamine metabolism is also suggested to be involved in cell fate, with identified roles in erythropoiesis and stem cell pluripotency^3,4^.

At the cellular level, glutamine is imported by solute carrier proteins (SLCs), where glutamine is consumed in both the cytoplasm and mitochondria by glutaminase. In the cytoplasm, glutamine is synthesized by glutamine synthetase from glutamate and ammonia. This subcellular compartmentalization is suggested to allow cells to precisely respond to metabolic need and partition the use of glutamine^5^. Despite having distinct compartments for glutamine metabolism, the dynamics of subcellular glutamine production and use remain poorly characterized. Towards illuminating glutamine metabolism, genetically encoded biosensors for glutamine have been developed^6,7^. The first glutamine biosensor was a Förster Resonance Energy Transfer (FRET)-based sensor, which was used to measure glutamine import and export in single cells^6^. A more recently developed circularly permuted yellow fluorescent protein (cpYFP)-based biosensor was used to illuminate differences in glutamine metabolism in the cytoplasm and mitochondrial matrix^7^. However, subtle glutamine dynamics across multiple compartments, cell types, and real-time crosstalk with other metabolic pathways has yet to be fully investigated.

To better illuminate the subcellular regulation of glutamine metabolism, we developed intracellular glutamine optical reporter (iGlo), a green fluorescent protein-based biosensor, to measure the subcellular dynamics of glutamine. Using iGlo, we measured subcellular glutamine uptake, production, and consumption, uncovering the real-time dynamics of glutamine metabolism in multiple cell types. Then, we performed multiplexed imaging of iGlo and a red-shifted lactate biosensor to investigate the temporal crosstalk between glutamine and glucose metabolism in single cells. Altogether, we present a new glutamine biosensor for an improved understanding of glutamine metabolism, uncovering the subcellular dynamics of this central metabolite.

## Results

### Development and characterization of iGlo

To design an intracellular glutamine optical reporter (iGlo), we selected the *Escherichia coli* periplasmic binding protein GlnH, part of the L-glutamine ATP-binding cassette transporter, as a sensing domain (Fig S1a)^8–10^. Consistent with the design of other biosensors, circularly permuted superfolder enhanced green fluorescent protein (cpsfEGFP), flanked by amino acid linkers, was inserted at four positions in the hinge region of GlnH, which undergoes conformational changes upon binding glutamine (Fig 1a)^11–14^. These four iGlo candidates were expressed in *E. coli* and screened in clarified lysate for changes in fluorescence upon glutamine addition. One candidate, with cpsfEGFP inserted after Ser 88 of GlnH, exhibited a small change in fluorescence upon glutamine addition (maximum fluorescence change, ΔF/F_0_ = (F_gln_ – F_PBS_)/F_PBS_; ΔF/F_0 480 nm_ = −0.12; Fig 1b). This candidate also exhibited a ratiometric fluorescence response when measuring both the 405 nm excitation-emission and 480 nm excitation-emission, whereas other candidates did not (R = F_405 nm_/F_480 nm_; ΔR/R_0_ = (R_gln_-R_PBS_)/R_PBS_; ΔR/R_0_ = 0.18). We selected this candidate for further optimization. To improve dynamic range, we performed two rounds of random mutagenesis targeting the linkers connecting cpsfEGFP to GlnH N- and C-termini and screened the resulting candidate biosensors in clarified *E. coli* lysate (Fig 1c). During screening, we evolved towards an intensiometric biosensor, prioritizing change in fluorescence (ΔF/F_0_) of 480 nm excitation-emission over ratiometric response (ΔR/R_0_), consistent with the design of other biosensors^12,15,16^. Top candidates from each round of linker screening, iGlo0.1 and iGlo0.2, were expressed in HeLa cells, and ΔF/F_0_ in response to glutamine measured (iGlo0.1 ΔF/F_0_ 1.52 ± 0.85; iGlo0.2 ΔF/F_0_ 3.72 ± 1.34; Fig 1d). iGlo0.2, which had a superior dynamic range, was selected as the final construct, and is subsequently referred to as “iGlo”.

**Figure 1.**
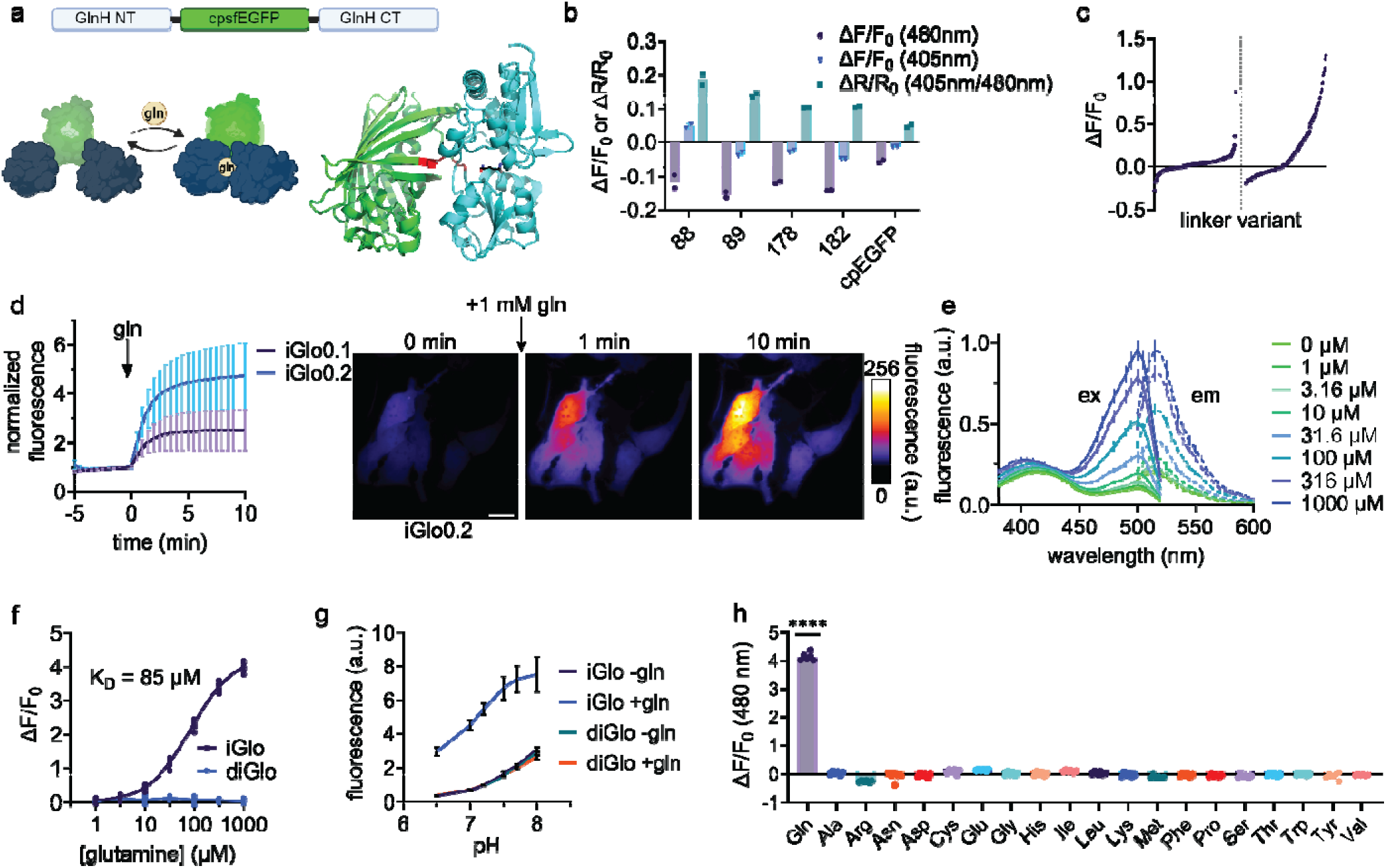
Development and characterization of iGlo. a, (top) The domain architecture of the iGlo biosensor design with cpsfEGFP inserted into GlnH. NT, N-termini; CT, C-termini. (bottom left) A cartoon representation of iGlo showing an increase in fluorescence upon reversibly binding glutamine. (bottom right) An AlphaFold3 model of iGlo (green, cpsfEGFP; red, linkers; cyan, GlnH; black, glutamine) b, Fractional changes of fluorescence and fluorescence ratio (ΔF/F_0_ and ΔR/R_0_) of initial candidate iGlo variants with cpsfEGFP inserted at four different sites in the hinge region of GlnH, as indicated by the residue number of the insertion site. c, ΔF/F_0_ from 480 nm excitation of candidate iGlo variants screened from site saturation mutagenesis libraries of the N-terminal linker (1st linker screen) and the C-terminal linker (2nd linker screen). d, Normalized fluorescence of iGlo0.1 (purple; n = 74 cells from 3 independent experiments) and iGlo0.2 (blue, n = 97 cells from 3 independent experiments), obtained from the first and second linker screens, respectively, in HeLa cells after the addition of 1 mM glutamine. Pseudocolor images of iGlo0.2 fluorescence over time in response to 1 mM glutamine in HeLa cells are shown. Cells were incubated with 5 mM alanine for 25 minutes prior to imaging. e, Excitation and emission spectra of iGlo from 0-1000 µM glutamine (n = 8 from 2 independent protein preparations). f, Maximum fluorescence response (ΔF/F_0_) of iGlo and diGlo in vitro to varying concentrations of glutamine (n = 8 from 2 independent protein preparations). K_D_ determined by nonlinear fit to Y=B_max_*X/(K_D_ + X). g, pH dependency of iGlo and diGlo in the presence of 0 mM (iGlo dark blue, diGlo teal; n = 8 from 2 individual protein preparations) or 1 mM glutamine (iGlo, light blue; diGlo orange; n = 8 from 2 individual protein preparations). h, ΔF/F_0_ of iGlo in response to 1 mM of each respective amino acid (n = 8 from 2 independent protein preparations). **** P < 0.0001 by ordinary one-way ANOVA. Scale bar represents 20 µm. For all graphs, mean ± standard deviation is shown.

Next, we characterized iGlo in vitro. iGlo exhibits an excitation peak at 500 nm, an emission peak at 515 nm, and a binding affinity for glutamine of 85 ± 6 µM (95% CI 79-91 µM) (Fig 1e-f). To generate a control biosensor, we introduced Arg75Met into GlnH, a mutation that has been reported to disrupt binding to glutamine^6,7^. This variant, termed “dead iGlo” or “diGlo”, was insensitive to glutamine (Fig 1f). We assessed the pH dependency of iGlo, as EGFP-based biosensors are known to be pH sensitive^17–19^. Across a range of physiologically relevant pH 6.5 – 8.0, iGlo increases in fluorescence with increasing pH, corresponding to decreasing ΔF/F_0_ as pH increases (Fig 1g and Fig S1b). Importantly, the pH dependency of diGlo closely matches that of glutamine-free iGlo. Therefore, we use diGlo as a control for all subsequent experiments. We confirmed iGlo is selective for glutamine over all other amino acids (ΔF/F_0_ = 4.14 ± 0.14; Fig 1h).

We utilized stopped flow to determine the in vitro kinetics of iGlo. We found that, like other biosensors, iGlo exhibits a dose-dependent increase in fluorescence that closely follows a two-phase association, with R^2^ values up to 0.9998 (Fig S1c)^20^. The resulting K_fast_ and K_slow_ values exhibit hyperbolic decay with respect to glutamine concentration (Fig S1d-e), potentially suggesting conformational selection binding mechanism^21,22^.

To demonstrate the utility of iGlo for multiple assays, we performed flow cytometry using iGlo fused to an mRuby3 expression marker (mRuby3-iGlo). We found iGlo reported an increase in fluorescence in HeLa cells treated with glutamine, as measured using flow cytometry (Fig S1f). Conversely, mRuby3-diGlo did not exhibit a similar change (Fig S1g). The increase in iGlo fluorescence in mRuby3-iGlo expressing cells corresponded with a significant increase in cpsfEGFP/mRuby3 fluorescence ratio, but not in mRuby3-diGlo expressing cells (iGlo cpsfEGFP/mRuby3 = 0.94 ± 0.011; diGlo cpsfEGFP/mRuby3 = 0.66 ± 0.007; Fig S1h). Thus, we have developed iGlo as a selective and sensitive glutamine biosensor, which, along with the control construct diGlo, can be used to measure intracellular glutamine dynamics in multiple assay formats.

### Measurement of subcellular glutamine import and export with iGlo

Glutamine is synthesized and consumed in distinct compartments, including the cytoplasm and mitochondrial matrix. Glutamine transporters on the plasma membrane enable the import of extracellular glutamine, and a putative mitochondrial glutamine transporter has been identified^23,24^. To investigate dynamics of glutamine import at distinct subcellular locations, we used targeting motifs to localize iGlo to the cytoplasm, mitochondrial matrix, and the nucleus as well as on the cytoplasmic faces of the outer mitochondrial, plasma, and lysosomal membranes and expressed each targeted iGlo in HeLa cells (Fig 2a). To measure maximal glutamine uptake at each compartment, cells expressing different localized iGlo variants were treated with alanine prior to imaging, which the sodium-dependent SLC1A5 obligatory exchanger transporter (ASCT2) imports while exporting glutamine^25^. When HeLa cells were treated with glutamine, we observed robust import of glutamine to all subcellular locations measured (iGlo ΔF/F_0_: cyto = 4.54 ± 1.15; matrix = 2.13 ± 0.32; nuc = 4.16 ± 0.82; OMM = 3.76 ± 0.93; PM = 4.44 ± 0.97; lyso = 2.35 ± 0.48; Fig 2b, Fig S2a-f). We next assessed iGlo’s ability to report glutamine uptake in multiple cell types, focusing on glutamine uptake into the cytoplasm and mitochondrial matrix, hot spots for glutamine metabolism. Cytoplasmic or mitochondrial matrix-targeted iGlo and corresponding diGlo variants were expressed in U2OS osteosarcoma cells or A549 non-small cell lung cancer cells. We observed robust glutamine uptake in the cytoplasm and matrix in both cell types (iGlo ΔF/F_0_: U2OS cyto = 5.32 ± 0.75; U2OS matrix = 1.96 ± 0.51; A549 cyto = 4.68 ± 0.34; A549 matrix = 1.53 ± 0.46; Fig 2c-d, Fig S2g-j).

**Figure 2.**
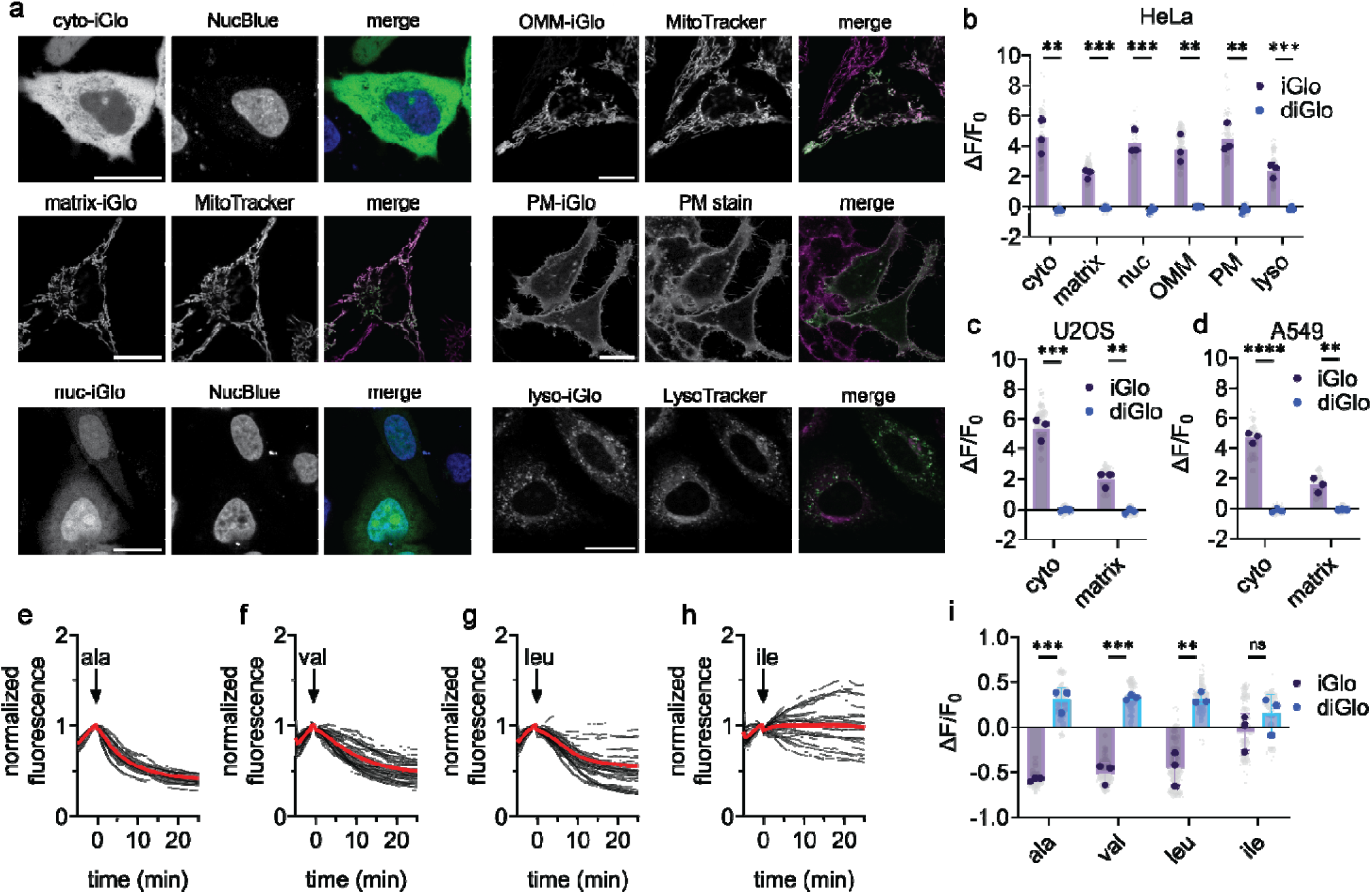
Measurement of subcellular glutamine import and export with iGlo. a, Representative images of HeLa cells expressing subcellular targeted iGlo to the cytoplasm, mitochondrial matrix, nucleus, outer mitochondrial membrane, plasma membrane, and lysosomes. Scale bar represents 20 µm. b, Fractional change in fluorescence (ΔF/F_0_) of HeLa cells expressing subcellular targeted iGlo or diGlo following the addition of 1 mM glutamine. Grey dots represent single cells. Solid dots (iGlo, purple; diGlo, blue) represent the mean of single cell measurements from biological replicates. Cyto ** P = 0.0020; matrix *** P = 0.0003; nuc *** P = 0.0008; OMM ** P = 0.0021; PM ** P = 0.0012; lyso *** P = 0.0009 by unpaired two-tailed t test. Cyto-iGlo n = 39 cells from 3 independent experiments. Cyto-diGlo n = 44 cells from 3 independent experiments. Matrix-iGlo n = 71 cells from 3 independent experiments. Matrix-diGlo n = 46 cells from 3 independent experiments. Nuc-iGlo n = 16 cells from 3 independent experiments. Nuc-diGlo = 19 cells from 3 independent experiments. OMM-iGlo n = 48 cells from 3 independent experiments. OMM-diGlo n = 30 cells from 3 independent experiments. PM-iGlo n = 21 cells from 3 independent experiments. PM-diGlo n = 15 cells from 3 independent experiments. Lyso-iGlo n = 28 cells from 3 independent experiments. Lyso-diGlo n = 25 cells from 3 independent experiments. c, ΔF/F_0_ of U2OS cells expressing cytoplasmic or mitochondria matrix targeted iGlo or diGlo following the addition of 1 mM glutamine. Grey dots represent single cell measurements. Solid dots (iGlo, purple; diGlo, blue) represent the mean of single cell measurements from biological replicates. Cyto *** P = 0.0002; matrix ** P = 0.0022 by unpaired two-tailed t test. Cyto-iGlo n = 46 cells from 3 independent experiments. Cyto-diGlo n = 34 cells from 3 independent experiments. Matrix-iGlo n = 33 cells from 3 independent experiments. Matrix-diGlo n = 43 cells from 3 independent experiments. d, ΔF/F_0_ of A549 cells expressing cytoplasmic or mitochondria matrix targeted iGlo or diGlo following the addition of 1 mM glutamine. Grey dots represent single cell measurements. Solid dots (iGlo, purple; diGlo, blue) represent the mean of single cell measurements from biological replicates. Cyto **** P < 0.0001; matrix ** P = 0.0036 by unpaired two-tailed t test. Cyto-iGlo n = 25 cells from 3 independent experiments. Cyto-diGlo n = 30 cells from 3 independent experiments. Matrix-iGlo n = 21 cells from 3 independent experiments. Matrix-diGlo n = 22 cells from 3 independent experiments. e-h, Mean (red) and single cell (black) normalized fluorescence of cyto-iGlo-expressing HeLa cells following the addition of 5 mM alanine (h, n = 46 cells), valine (i, n = 69 cells), leucine (j, n = 69 cells), and isoleucine (k, n = 53 cells) across three biological replicates. i, ΔF/F_0_ of HeLa cells expressing cyto-iGlo (purple) or cyto-diGlo (blue) following the addition of 5 mM alanine, isoleucine, leucine, or valine. Grey dots represent single cell measurements. Solid dots represent the mean of single cell measurements from biological replicates. Ala *** P = 0.0003; val *** P = 0.0003; leu ** P = 0.0025; ile n.s. P = 0.29 by unpaired two-tailed t test. Ala cyto-iGlo n = 46 cells from 3 independent experiments. Ala cyto-diGlo n = 47 cells from 3 independent experiments. Val cyto-iGlo n = 69 cells from 3 independent experiments. Val cyto-diGlo n = 44 cells from 3 independent experiments. Leu cyto-iGlo n = 69 cells from 3 independent experiments. Leu cyto-diGlo n = 46 cells from 3 independent experiments. Ile cyto-iGlo n = 53 cells from 3 independent experiments. Ile cyto-diGlo n = 34 cells from 3 independent experiments.

In addition to being exchanged with alanine through SLC1A5, glutamine is exchanged for branched-chain amino acids (BCAAs) through SLC7A5 (LAT1)^26^. We sought to use iGlo to determine if glutamine export from the cytoplasm would vary based on the amino acid used to exchange glutamine. We found that in HeLa cells, alanine and valine treatment resulted in efficient export of glutamine (iGlo ΔF/F_0_: ala = −0.58 ± 0.02; val = −0.51 ± 0.12; Fig 2e-f, i). Meanwhile, leucine resulted in overall depletion of cytoplasmic glutamine, with large cell-to-cell heterogeneity, and isoleucine was not efficient in inducing glutamine export compared to diGlo control (iGlo ΔF/F_0_: leu = −0.45 ± 0.19; ile = −0.05 ± 0.20; Fig 2g-i). Thus, using iGlo, we find that glutamine is transported throughout the cell, and the efflux of glutamine through known SLCs is dependent on the amino acid used to exchange with glutamine.

As alanine-mediated export of glutamine was efficient, we then used alanine to determine compartment-specific import and export dynamics of glutamine. In HeLa, U2OS, and A549 cells expressing either cytoplasmic or mitochondrial matrix-targeted iGlo, we measured glutamine import into the cytoplasm and mitochondrial matrix, which was reversed with alanine (Fig S2k-p). As there is no known alanine-glutamine exchanger on the mitochondrial membrane, these data suggest that the cytoplasmic and mitochondrial pools of glutamine might be closely linked^27^.

### iGlo reveals real-time cell type-specific dynamics of glutamine metabolism

The compartmentalized regulation of glutamine metabolism has been suggested to provide a mechanism for cells to rapidly adjust to changing metabolic need^5^. We hypothesized that perturbing glutamine metabolism would have distinct subcellular impacts on glutamine pools. To test our hypothesis, we focused on glutamine dynamics in the cytoplasm and mitochondrial matrix of HeLa, U2OS, and A549 cells, all of which are reported to use glutamine to fuel their metabolism or to be glutamine dependent^28–30^. First, we targeted glutamine synthetase, which produces glutamine through the ATP-dependent condensation of glutamate and ammonia^31^. To measure the dynamics of glutamine synthesis, we treated cells expressing iGlo targeted to the cytoplasm or mitochondrial matrix with ammonium chloride (NH_4_Cl). We found HeLa cells readily and rapidly produce glutamine from ammonia, which we observed as an increase in iGlo fluorescence in the cytoplasm and mitochondrial matrix compared to diGlo control (iGlo ΔF/F_0_ in HeLa: cyto = 1.03 ± 0.13; matrix = 0.44 ± 0.19; Fig 3a-d). Pretreatment of HeLa cells with a glutamine synthetase inhibitor, methionine sulfoximine (MSO), abolished the NH_4_Cl-dependent increase in iGlo fluorescence in both the cytoplasm and mitochondrial matrix (iGlo ΔF/F_0_ in HeLa with MSO pretreatement: cyto = −0.20 ± 0.03; matrix = −0.18 ± 0.03; Fig S3a-b)^32^. However, U2OS and A549 cells did not significantly increase glutamine with ammonia treatment as compared to diGlo control (iGlo ΔF/F_0_: U2OS cyto = −0.02 ± 0.13; A549 cyto = 0.03 ± 0.07; U2OS matrix = 0.03 ± 0.09; A549 matrix = 0.03 ± 0.07; Fig 3c-d). To confirm iGlo is accurately reporting glutamine production, we measured glutamine production in each cell line treated with NH_4_Cl using LC-MS metabolomics, which was consistent with our imaging results (normalized glutamine ion counts: HeLa = 2.29 ± 0.45; U2OS = 0.83 ± 0.17; A549 = 1.17 ± 0.18; Fig S3c).

**Figure 3.**
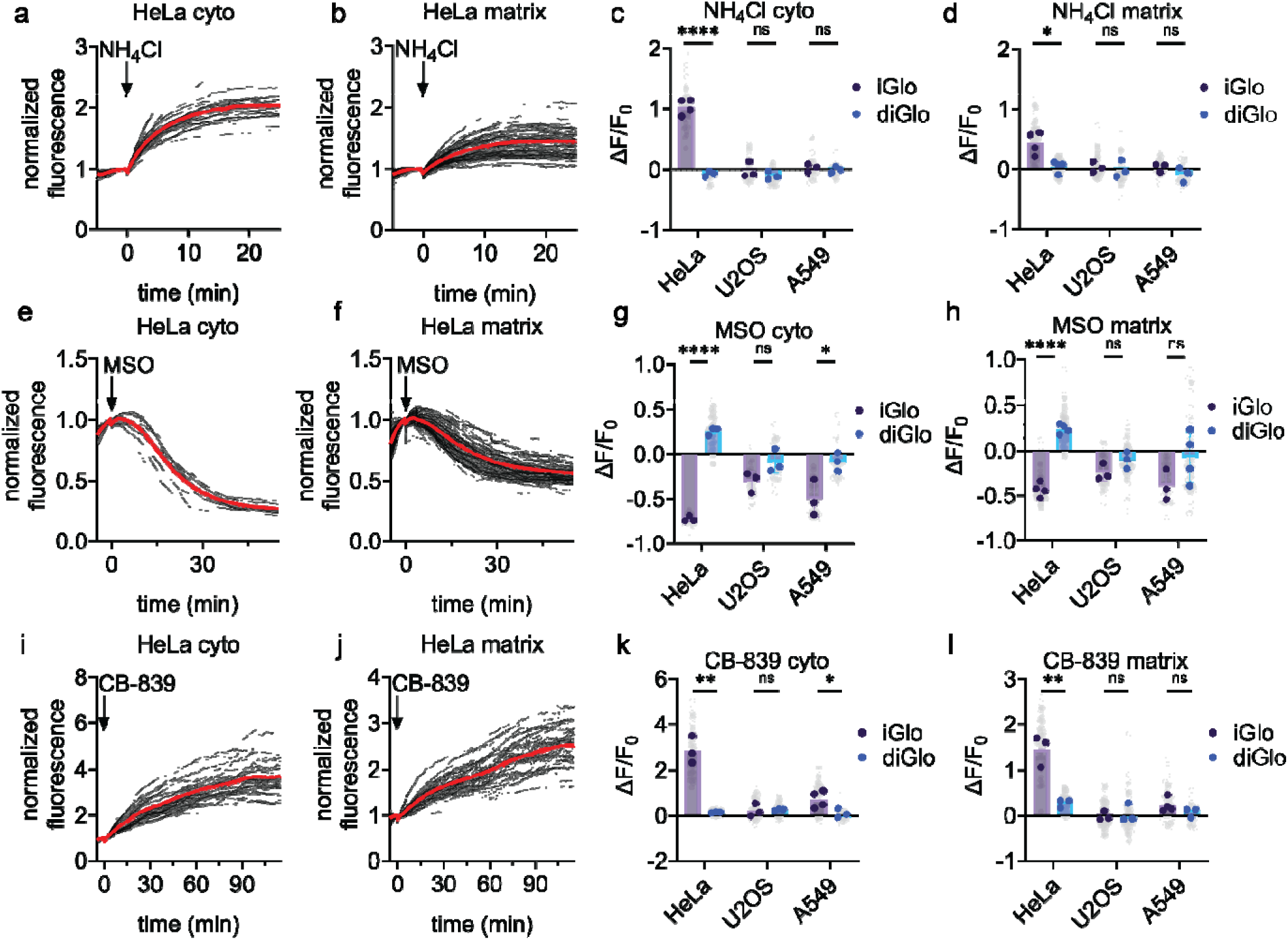
iGlo reveals real-time dynamics of glutamine metabolism. a-b, Mean (red) and single cell (black) normalized fluorescence of cyto-iGlo (a, n = 46 cells) and matrix-iGlo (b, n = 77 cells) in HeLa cells following the addition of 200 µM NH_4_Cl across four biological replicates. Cells were treated with 5 mM alanine for 25 minutes prior to imaging. c, Fractional change in fluorescence (ΔF/F_0_) of HeLa, U2OS, and A549 cells expressing cyto-iGlo or cyto-diGlo following the addition of 200 µM NH_4_Cl. HeLa **** P < 0.0001; U2OS n.s. P = 0.37; A549 n.s. P = 0.61 by unpaired two-tailed t test. HeLa cyto-iGlo n = 46 cells from 4 independent experiments. HeLa cyto-diGlo n = 38 cells from 3 independent experiments. U2OS cyto-iGlo n = 38 cells from 3 independent experiments. U2OS cyto-diGlo n = 35 cells from 3 independent experiments. A549 cyto-iGlo n = 24 cells from 3 independent experiments. A549 cyto-diGlo n = 20 cells from 3 independent experiments. d, ΔF/F_0_ of HeLa, U2OS, and A549 cells expressing matrix-iGlo or matrix-diGlo following the addition of 200 µM NH_4_Cl. HeLa * P = 0.010; U2OS n.s. P = 0.74; A549 n.s. P = 0.23 by unpaired two-tailed t test. HeLa matrix-iGlo n = 77 cells from 4 independent experiments. HeLa matrix-diGlo n = 86 cells from 4 independent experiments. U2OS matrix-iGlo n = 62 cells from 3 independent experiments. U2OS matrix-diGlo n = 45 cells from 3 independent experiments. A549 matrix-iGlo n = 28 cells from 3 independent experiments. A549 matrix-diGlo n = 26 cells from 3 independent experiments. e-f, Mean (red) and single cell (black) normalized fluorescence of cyto-iGlo (e, n = 24 cells) and matrix-iGlo (f, n = 89 cells) in HeLa cells following the addition of 5 mM MSO across three and four biological replicates, respectively. g, ΔF/F_0_ of HeLa, U2OS, and A549 cells expressing cyto-iGlo or cyto-diGlo following the addition of 5 mM MSO. HeLa **** P < 0.0001; U2OS n.s. P = 0.095; A549 * P = 0.033 by unpaired two-tailed t test. HeLa cyto-iGlo n = 22 cells from 3 independent experiments. HeLa cyto-diGlo n = 49 cells from 3 independent experiments. U2OS cyto-iGlo n = 38 cells from 3 independent experiments. U2OS cyto-diGlo n = 34 cells from 3 independent experiments. A549 cyto-iGlo n = 33 cells from 3 independent experiments. A549 cyto-diGlo n = 26 cells from 3 independent experiments. h, ΔF/F_0_ of HeLa, U2OS, and A549 cells expressing matrix-iGlo or matrix-diGlo following the addition of 5 mM MSO. HeLa **** P < 0.0001; U2OS n.s. P = 0.13; A549 n.s. P = 0.14 by unpaired two-tailed t test. HeLa matrix-iGlo n = 89 cells from 4 independent experiments. HeLa matrix-diGlo n = 101 cells from 4 independent experiments. U2OS matrix-iGlo n = 45 cells from 3 independent experiments. U2OS matrix-diGlo n = 52 cells from 3 independent experiments. A549 matrix-iGlo n = 31 cells from 3 independent experiments. A549 matrix-diGlo n = 47 cells from 3 independent experiments. i-j, Mean (red) and single cell (black) normalized fluorescence of cyto-iGlo (I, n = 55 cells) and matrix-iGlo (j, n = 57) in HeLa cells following the addition of 10 µM CB-839 across three biological replicates. Cells were treated with 5 mM alanine for 25 minutes prior to imaging. k, ΔF/F_0_ of HeLa, U2OS, and A549 cells expressing cyto-iGlo or cyto-diGlo following the addition of 10 µM CB-839. HeLa ** P = 0.0013; U2OS n.s. P = 0.86; A549 * P = 0.049 by unpaired two-tailed t test. HeLa cyto-iGlo n = 55 cells from 3 independent experiments. HeLa cyto-diGlo n = 37 cells from 3 independent experiments. U2OS cyto-iGlo n = 38 cells from 3 independent experiments. U2OS cyto-diGlo n = 38 cells from 3 independent experiments. A549 cyto-iGlo n = 54 cells from 4 independent experiments. A549 cyto-diGlo n = 18 cells from 3 independent experiments. l, ΔF/F_0_ of HeLa, U2OS, and A549 cells expressing matrix-iGlo or matrix-diGlo following the addition of 10 µM CB-839. HeLa ** P = 0.0043; U2OS n.s. P = 0.69; A549 n.s. P = 0.20 by unpaired two-tailed t test. HeLa matrix-iGlo n = 57 cells from 3 independent experiments. HeLa matrix-diGlo n = 61 cells from 3 independent experiments. U2OS matrix-iGlo n = 64 cells from 3 independent experiments. U2OS matrix-diGlo n = 56 cells from 3 independent experiments. A549 matrix-iGlo n = 41 cells from 4 independent experiments. A549 matrix-diGlo n = 24 cells from 3 independent experiments. Grey dots represent single cell measurements. Solid dots (iGlo, purple; diGlo, blue) represent the mean of single cell measurements from biological replicates.

Next, we used MSO to further probe the dynamics of glutamine production in these cell lines. In HeLa cells, we measured a decrease in glutamine in the cytoplasm and mitochondrial matrix compared to diGlo control (iGlo ΔF/F_0_ in HeLa: cyto = −0.73 ± 0.03; matrix = −0.44 ± 0.08; Fig 3e-h). In U2OS cells, MSO did not significantly alter glutamine dynamics in comparison to diGlo control (iGlo ΔF/F_0_ in U2OS: cyto = −0.31 ± 0.11; matrix = −0.25 ± 0.09; Fig 3g-h). A549 cells exhibited a significant decrease in glutamine after MSO treatment in the cytoplasm, but not mitochondrial matrix, compared to diGlo control (iGlo ΔF/F_0_ in A549: cyto = −0.50 ± 0.20; matrix = −0.40 ± 0.17; Fig 3g-h). We compared our findings with LC-MS metabolomics, finding that only HeLa cells exhibited a significant decrease in bulk glutamine abundance following MSO treatment compared to vehicle control (normalized glutamine ion counts: HeLa = 0.34 ± 0.06; U2OS = 1.19 ± 0.41; A549 = 0.83 ± 0.08; Fig S3d).

Finally, we targeted glutamine consumption by glutaminase, which hydrolyzes glutamine to form glutamate and ammonia^33^. HeLa, U2OS, and A549 cells expressing either cytoplasmic or mitochondrial matrix iGlo were treated with CB-839, a potent glutaminase inhibitor^34^. In HeLa cells, we observed an increase in glutamine in both the cytoplasm and mitochondrial matrix compared to diGlo (iGlo ΔF/F_0_ in HeLa: cyto = 2.84 ± 0.58; matrix = 1.45 ± 0.34; Fig 3i-l). In A549 cells, we observed a small increase in glutamine levels in the cytoplasm, but not the mitochondrial matrix, as compared to diGlo control (iGlo ΔF/F_0_ in A549: cyto = 0.71 ± 0.37; matrix = 0.22 ± 0.16; Fig 3k-l). However, U2OS cells did not have a significant change in glutamine levels in either the cytoplasm or mitochondrial matrix (iGlo ΔF/F_0_ in U2OS: cyto = 0.22 ± 0.28; matrix = −0.02 ± 0.10; Fig 3k-l). We supported these findings using LC-MS metabolomics, although, unlike with cyto-iGlo, the measured increase in glutamine in A549 was not statistically significant (normalized glutamine ion counts: HeLa = 5.41 ± 0.70; U2OS = 1.11 ± 0.06; A549 = 1.26 ± 0.11; Fig S3e).

### Culture conditions impact glutamine metabolism in HepG2 cells

Building off our work with HeLa, U2OS, and A549 cells, we sought to study the dynamics of glutamine metabolism in HepG2 cells, which have been reported to use glutamine for anaplerosis and are glutamine dependent^35^. However, when we expressed iGlo in the cytoplasm of HepG2 cells, we observed large cell-to-cell variability in glutamine uptake, with overall very low levels of glutamine import, unlike other cell lines tested (iGlo ΔF/F_0_ = 0.84 ± 0.23; Fig 4a). We hypothesized that the culture medium HepG2 cells were grown in (Dulbecco’s Modified Eagle Medium, DMEM), which contains non-physiologically relevant concentrations of glucose and amino acids, might be providing sufficient nutrients for HepG2 growth, limiting uptake of more exogenous glutamine. To test this hypothesis, we instead cultured HepG2 cells in human plasma-like medium (HPLM), which contains physiological concentrations of glutamine, and measured glutamine import (see Methods for concentration information). After culturing in HPLM, we observed increased glutamine uptake, as compared to diGlo and cells cultured in DMEM (iGlo ΔF/F_0_ = 2.18 ± 1.39; Fig 4b-c, Fig S4a).

**Figure 4.**
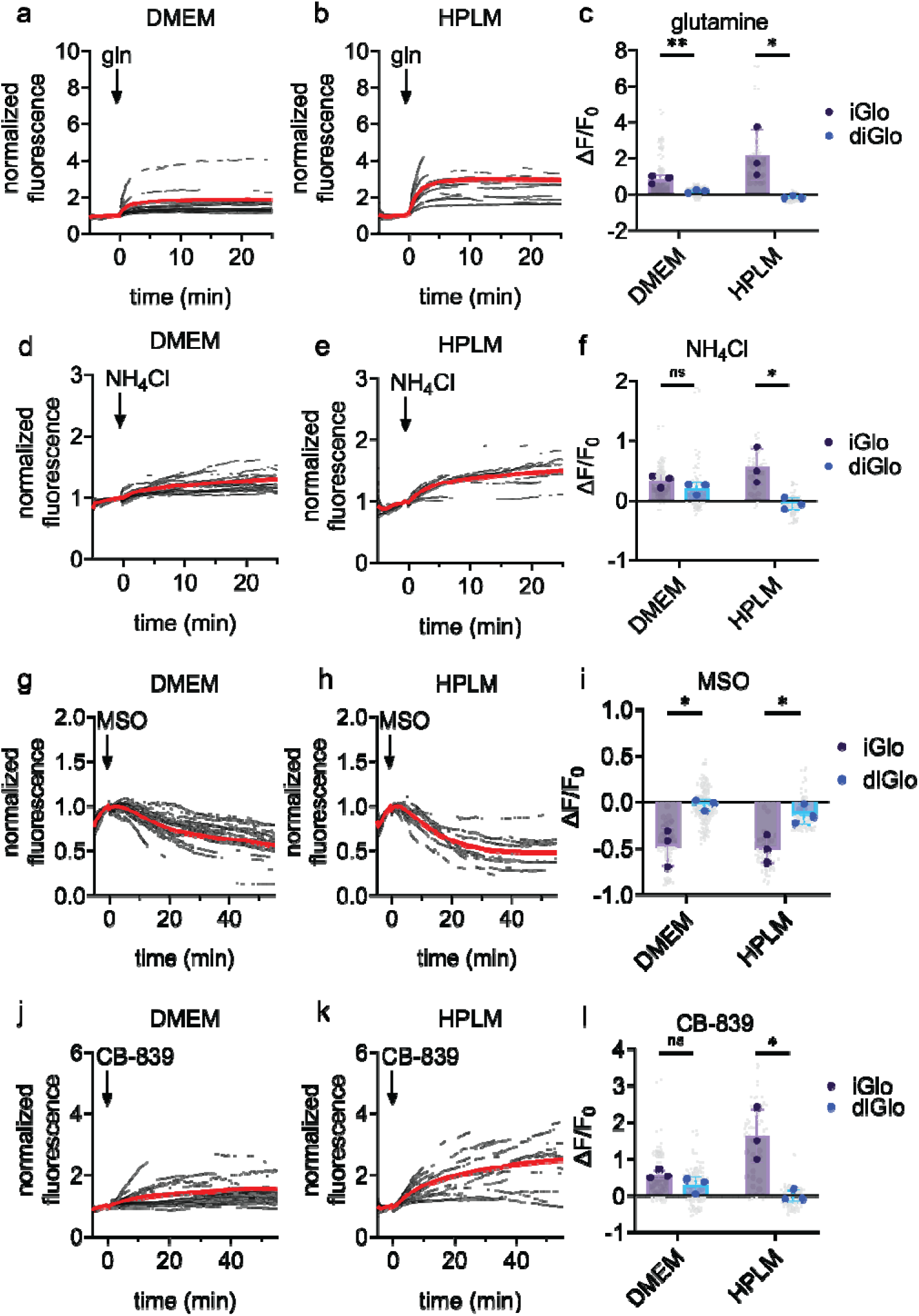
Culture conditions impact glutamine metabolism in HepG2 cells. a-b, Mean (red) and single cell (black) normalized fluorescence of cyto-iGlo in HepG2 cells either cultured in DMEM only (a, n = 38 cells) or exchanged for HPLM 24 hours prior to imaging (b, n = 29) treated with 1 mM glutamine across three biological replicates. Cells were treated with 5 mM alanine for 25 minutes prior to imaging. c, ΔF/F_0_ of HepG2 cells expressing cyto-iGlo (purple) or cyto-diGlo (blue) 25 minutes after the addition of 1 mM glutamine. DMEM n.s. P = 0.11; HPLM * P = 0.019 by unpaired two-tailed t test. DMEM iGlo n = 38 from 3 independent experiments. DMEM diGlo = 32 cells from 3 independent experiments. HPLM iGlo n = 29 cells from 3 independent experiments. HPLM diGlo n = 24 cells from 3 independent experiments. d-e, Mean (red) and single cell (black) normalized fluorescence of cyto-iGlo in HepG2 cells either cultured in DMEM only (d, n = 37 cells) or exchanged for HPLM 24 hours prior to imaging (e, n = 26 cells) treated with 200 µM NH_4_Cl across three biological replicates. Cells were treated with 5 mM alanine for 25 minutes prior to imaging. f, Fractional change in fluorescence (ΔF/F_0_) of live HepG2 cells expressing cyto-iGlo (purple) or cyto-diGlo (blue) 25 minutes after the addition of 200 µM NH_4_Cl. DMEM ** P = 0.0089; HPLM * P = 0.045 by unpaired two-tailed t test. DMEM iGlo n = 37 from 3 independent experiments. DMEM diGlo = 31 cells from 3 independent experiments. HPLM iGlo n = 26 cells from 3 independent experiments. HPLM diGlo n = 26 cells from 3 independent experiments. g-h, Mean (red) and single cell (black) normalized fluorescence of cyto-iGlo in HepG2 cells either cultured in DMEM only (h, n = 43 cells) or exchanged for HPLM 24 hours prior to imaging (i, n = 24 cells) treated with 5 mM MSO across three biological replicates. i, ΔF/F_0_ of HepG2 cells expressing cyto-iGlo (purple) or cyto-diGlo (blue) 55 minutes after the addition of 5 mM MSO. DMEM n.s. P = 0.22; HPLM * P = 0.029 by unpaired two-tailed t test. DMEM iGlo n = 43 from 3 independent experiments. DMEM diGlo = 47 cells from 3 independent experiments. HPLM iGlo n = 24 cells from 3 independent experiments. HPLM diGlo n = 23 cells from 3 independent experiments. j-k, Mean (red) and single cell (black) normalized fluorescence of cyto-iGlo in HepG2 cells either cultured in DMEM only (l, n = 41 cells) or exchanged for HPLM 24 hours prior to imaging (m, n = 32 cells) treated with 10 µM CB-839 across three biological replicates. Cells were treated with 5 mM alanine for 25 minutes prior to imaging. l, ΔF/F_0_ of HepG2 cells expressing cyto-iGlo (purple) or cyto-diGlo (blue) 55 minutes after the addition of 10 µM CB-839. DMEM * P = 0.021; HPLM * P = 0.024 by unpaired two-tailed t test. DMEM iGlo n = 41 from 3 independent experiments. DMEM diGlo = 41 cells from 3 independent experiments. HPLM iGlo n = 32 cells from 3 independent experiments. HPLM diGlo n = 22 cells from 3 independent experiments.

We next hypothesized that the cell culture conditions HepG2 were cultured in would have an impact on glutamine utilization. To test this hypothesis, we treated HepG2 cells expressing iGlo grown in either DMEM or HPLM with NH_4_Cl, finding that culturing in HPLM significantly enhanced the number of cells producing glutamine and the magnitude of glutamine production as compared to diGlo control (iGlo DMEM ΔF/F_0_ = 0.33 ± 0.11; HPLM ΔF/F_0_ = 0.56 ± 0.30; Fig 4d-f, Fig S4b). We confirmed this result using LC-MS metabolomics (normalized glutamine ion counts: DMEM = 0.88 ± 0.08; HPLM = 2.02 ± 0.56; Fig S4c). We next treated HepG2 cells expressing iGlo with MSO, finding that inhibition of glutamine synthetase resulted in similar decreases in glutamine levels in cells grown in either DMEM or HPLM, confirmed using metabolomics (iGlo DMEM ΔF/F_0_ = −0.48 ± 0.20; HPLM ΔF/F_0_ = −0.51 ± 0.14; Fig 4g-i, Fig S4d). We supported this finding using LC-MS metabolomics (normalized glutamine ion counts: DMEM = 0.28 ± 0.03; HPLM = 0.46 ± 0.09; Fig S4e). Finally, we treated HepG2 cells expressing iGlo with CB-839 and found that inhibition of glutaminase substantially increased accumulation of glutamine in single cells and overall glutamine levels in cells grown in HPLM (iGlo DMEM ΔF/F_0_ = 0.58 ± 0.13; HPLM ΔF/F_0_ = 1.63 ± 0.72; Fig 4j-l, Fig S4f). We supported this result with LC-MS metabolomics (normalized glutamine ion counts: DMEM = 1.84 ± 0.40; HPLM = 6.24 ± 0.44; Fig S4g). We found that more cells were increasing glutaminolysis when cultured in HPLM compared to DMEM. Thus, we find that at least for HepG2 cells, media conditions significantly impact the dynamics of their glutamine metabolism, reflected in changes in shifts in glutamine metabolism at the single cell level as measured with iGlo.

### Multiplexed imaging with iGlo reveals rapid metabolic adaptations

Glutamine can be used to provide carbons for the tricarboxylic acid cycle (TCA cycle) for both oxidation and macromolecular biosynthesis, particularly during cell proliferation. The TCA cycle is canonically fueled by glucose, but many cancer cell types, including HeLa cells, reprogram the cycle to rely on glutamine, and direct glucose towards glycolysis and lactate production^28,36^. We hypothesized that when glutaminolysis is inhibited with CB-839, HeLa cells would increase glycolysis to compensate for the inhibition of glutamine-derived carbon flux into the TCA cycle, thus increasing lactate production. To test this hypothesis, we performed multiplexed imaging of HeLa cells expressing both iGlo and a red lactate biosensor, R-iLACCO1.2 (Fig 5a)^37^. These cells were treated with CB-839, and dynamics of both glutamine and lactate measured. We found that when glutamine incorporation into the TCA cycle was impaired, both glutamine and lactate levels increased in single cells (iGlo ΔF/F_0_ = 3.15 ± 0.63; R-iLACCO1.2 ΔF/F_0_ = 0.51 ± 0.09; Fig 5b-c, Fig S5a). Under these conditions, lactate accumulation was rapid, occurring faster than glutamine accumulation (iGlo t_1/2_ = 29.0 ± 8.6 min; R-iLACCO1.2 t_1/2_ = 12.0 ± 2.3 min; Fig 5d), suggesting a quick switch in metabolism occurs when glutamine consumption is inhibited.

**Figure 5.**
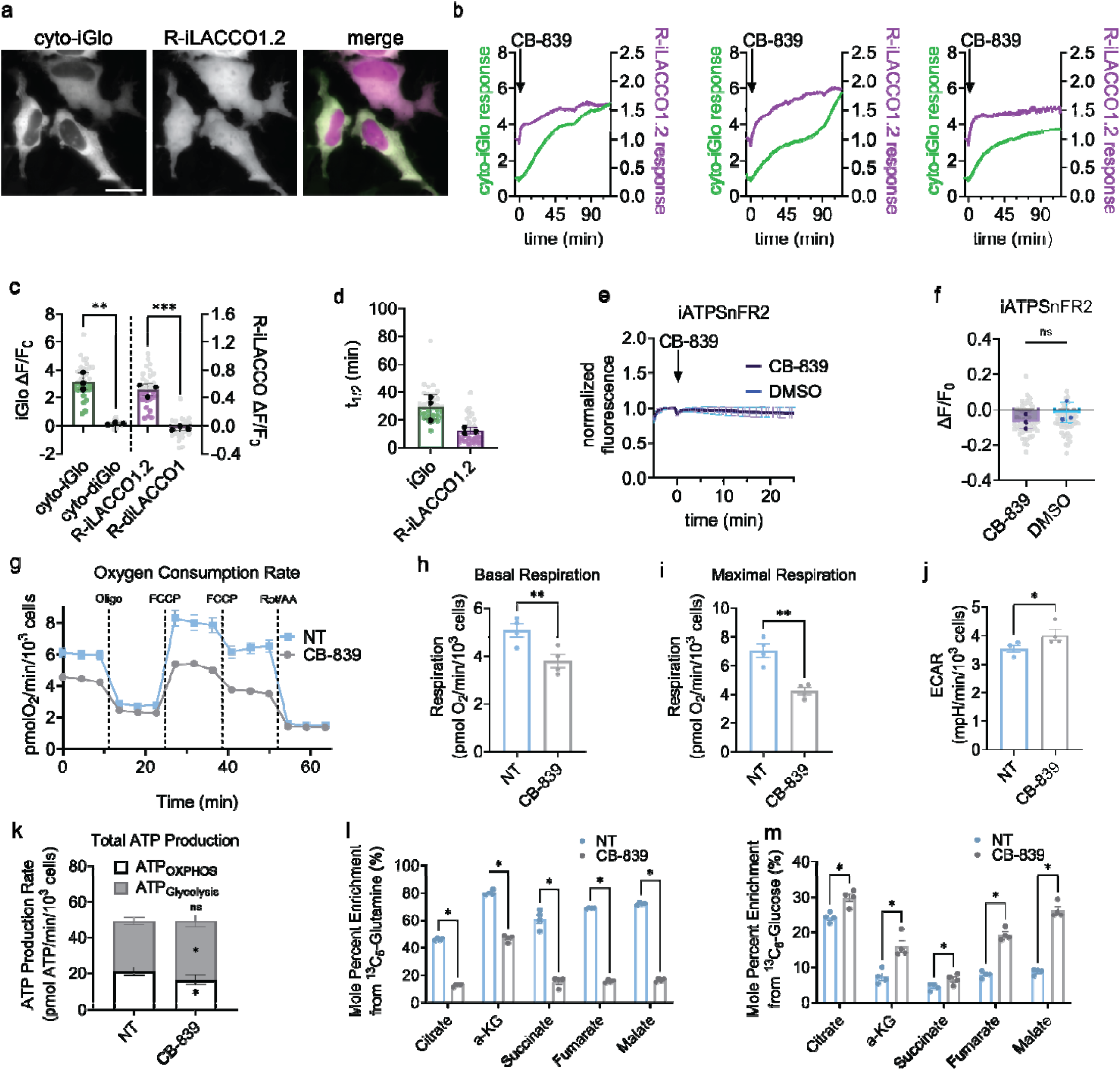
Multiplexed imaging with iGlo reveals rapid metabolic adaptations. a, Representative HeLa cells expressing cyto-iGlo (green) and R-iLACCO1.2 (magenta). Scale bar represents 20 µm. b, Representative single cell traces from each biological replicate of HeLa cells co-expressing cyto-iGlo (green) and R-iLACCO1.2 (magenta) treated with 10 µM CB-839. Y-axis labels refer to cyto-iGlo and R-iLACCO1.2 fluorescence of single cells normalized to the time point immediately preceding CB-839 addition. Cells were treated with 5 mM alanine prior to imaging for 25 minutes. c, Fractional change of fluorescence (ΔF/F_0_) of HeLa cells expressing cyto-iGlo and R-iLACCO1.2 or cyto-iGlo and R-diLACCO1 115 minutes after the addition of 10 µM CB-839. Grey dots represent single cells. cyto-iGlo v cyto-diGlo ** P = 0.0013; R-iLACCO1.2 v R-diLACCO1 *** P = 0.0007. Cyto-iGlo and R-iLACCO1.2 n = 32 cells from 3 independent experiments. Cyto-diGlo and R=diLACCO1 n = 23 cells from 3 independent experiments. d, Time to half maximum (t_1/2_) of single cells (grey dots) and biological replicates (black dots) of cyto-iGlo (green) and R-iLACCO1.2 (magenta) in HeLa cells following addition of 10 µM CB-839. e, Normalized fluorescence of HeLa cells expressing iATPSnFR2 treated with CB-839 (purple, n = 49 cells) or DMSO (blue, n = 51 cells). Traces represent mean ± S.D. of single cell measurements across three biological replicates. Cells were treated with 5 mM alanine prior to imaging. f, ΔF/F_0_ of HeLa cells expressing iATPSnFR2 after 25-minute treatments with CB-839 (purple, n = 49 cells) or DMSO (blue, n = 51 cells). Grey dots represent single cell measurements and solid dots represent the means of the three biological replicates. n.s. P = 0.28 by unpaired two-tailed t test. g, Oxygen consumption rate versus time of HeLa cells following treatment with 10 µM CB-839 or no treatment. h, Basal respiration of HeLa cells following treatment with 10 µM CB-839 or no treatment. ** P = 0.0011 by paired two-tailed t test. i, Maximal respiration of HeLa cells following treatment with 10 µM CB-839 or no treatment. ** P = 0.0010 by paired two-tailed t test. j, Extracellular acidification rate of HeLa cells following treatment with 10 µM CB-839 or no treatment. * P = 0.039 by paired two-tailed t test. k, ATP production from glycolysis and OXPHOS in HeLa cells following treatment with 10 µM CB-839 or no treatment. OXPHOS * P = 0.011; Glycolysis * P = 0.028; Total ATP n.s. P = 0.90 by paired two-tailed t test. l, Mole percent enrichment of citrate, α-ketoglutarate, succinate, fumarate, and malate from ^13^C_5_-glutamine in HeLa following 24-hour treatment with 2 µM CB-839 or no treatment. * P = 0.000004, 0.00013, 0.00014, 0.000003, and 0.000002, respectively, by paired t test using the Holm-Šídák method. m, Mole percent enrichment of citrate, α-ketoglutarate, succinate, fumarate, and malate from ^13^C_6_-glucose in HeLa following 24-hour treatment with 2 µM CB-839 or no treatment. * P = 0.0015, 0.0033, 0.017, 0.00050, 0.00033 by paired t test using the Holm-Šídák method.

We hypothesized that this rapid switch in metabolism could be to maintain constant ATP levels in cells to maintain basic cellular functions. To test this hypothesis, we expressed an ATP biosensor, iATPSnFR2, in HeLa cells^12^. We found that CB-839 treatment did not significantly alter ATP levels, compared to vehicle control (CB-839 ΔF/F_0_ = −0.07 ± 0.04; DMSO ΔF/F_0_ = - 0.02 ± 0.06; Fig 5e-f). As a control, we observed that iATPSnFR2 reports a decrease in ATP upon the inhibition of glycolysis under identical conditions when cells are treated with 2-deoxy-D-glucose (2DG, ΔF/F0 = −0.19 ± 0.03; DMSO ΔF/F0 = 0.00 ± 0.05; Fig S5b-c. Together, these data suggest that when glutamine incorporation into the TCA cycle is limited, HeLa cells rapidly upregulate glycolysis to prevent a decrease in ATP levels and maintain energy homeostasis.

To support our finding that HeLa cells become more glycolytic and less reliant on oxidative phosphorylation when glutaminolysis is inhibited, we measured the oxygen consumption rate and extracellular acidification rate of HeLa cells treated with CB-839. We found that respiration was decreased in CB-839-treated cells and the extracellular acidification rate was elevated (Fig 5g-j, Fig S5b). Consistent with our imaging data, total ATP production rate was comparable between non-treated and CB-839-treated cells, but under CB-839 treatment, glycolysis contributed more to total ATP production than oxidative phosphorylation (Fig 5k, Fig S5c-d). To further support our measurements made using iGlo and R-iLACCO1.2, we performed stable isotope tracing using ^13^C_6_-glucose or ^13^C_5_-glutamine. When HeLa cells were treated with CB-839, we measured decreased glutamine incorporation into the TCA cycle with a compensatory increase in glucose incorporation (Fig 5l-m). We also observed an increased lactate-to-pyruvate ratio, a metric reflecting increased glycolysis and independent of confounding effects from long-term CB-839 treatment on cell proliferation (Fig S5e). Thus, by co-imaging iGlo with the lactate biosensor R-iLACCO1.2 at the single-cell level, alongside measurements of respiration and stable isotope tracing, we identified a mechanism by which HeLa cells maintain stable ATP levels following inhibition of glutaminolysis by a rapid switch from oxidative phosphorylation to glycolysis.

## Discussion

Genetically encoded biosensors are powerful protein-based tools that enable the study of metabolic dynamics with high spatiotemporal resolution. Inspired by the importance of glutamine metabolism in cell function, we developed a EGFP-based intracellular glutamine optical reporter, iGlo. We then demonstrated the use of iGlo in multiple cell types, subcellular locations, and for multiplexed imaging, effectively illuminating the dynamics of glutamine in single cells. Additionally, we further validated the use of iGlo for flow cytometry-based assays, highlighting the versatility of iGlo. Biosensors have previously been developed for glutamine. A FRET-based glutamine reporter had a limited dynamic range of approximately 18%, hindering the broad applicability of the sensor^6^. Recently, a single fluorescent protein-based glutamine biosensor, GlutaR, has been developed, which is similar in conceptual design to iGlo, but incorporates distinct linker sequences, an alternative fluorescent protein, and minor modifications to the GlnH protein^7^. In comparison to GlutaR, iGlo exhibits a higher affinity with a K_D_ for glutamine of 85 µM, suggesting that iGlo may be more useful for imaging in low glutamine conditions^38,39^. Future work will focus on improving the functional properties of iGlo, including K_D_, K_on_, and dynamic range for use in a wider range of experimental and physiologically-relevant conditions.

With iGlo, we developed more subcellular targeted variants, not only to the cytoplasm, mitochondrial matrix, and nucleus, but also to the cytoplasmic face of the plasma membrane, outer mitochondrial membrane, and lysosomal membrane. We anticipate that having a wider array of subcellular targeted biosensors will facilitate future studies of compartmentalized glutamine metabolism. Altogether, we have added a new tool to the toolbox of biosensors to measure subcellular glutamine dynamics, enabling robust measurement of glutamine metabolism.

With iGlo, we demonstrated that intracellular glutamine can be depleted in the presence of exogenous amino acids, as SLC1A5 is reported to exchange alanine for glutamine and SLC7A5 is reported to exchange BCAAs for glutamine^40^. We find that the identity of the amino acid used to trigger glutamine export from HeLa cells significantly matters, as alanine, valine, and leucine were much more potent in depleting intracellular glutamine than isoleucine when measured at the single-cell level. Additionally, we found that alanine was able to deplete both the cytoplasm and mitochondrial matrix of glutamine. This would suggest that cytoplasmic and mitochondrial glutamine pools are tightly coupled, or a glutamine-alanine antiporter is present on the mitochondria. As glutamine transporters at the mitochondria are not well understood, future investigations could focus on the mechanisms of crosstalk or compartmentalization of glutamine pools^24^.

To further study how glutamine metabolism is subcellularly regulated, we investigated the impact of perturbing glutamine metabolism on cytoplasmic and mitochondrial matrix pools of glutamine in multiple cell types, including HeLa, U2OS, and A549 cells. Through our investigations, we uncovered both cell-to-cell variability and cell type-specific differences in glutamine dynamics following the inhibition of glutamine synthetase and glutaminase. For instance, HeLa cells were most efficient at incorporating ammonia into glutamine compared to U2OS and A549, with cell-to-cell variability in the dynamics of glutamine production observed within HeLa cells. Within an unsynchronized cell population, this variability could be due to differences in cell state and metabolic need, as seen with other readouts of cellular metabolism and energy sensing^37,41^. Further work into correlating cell cycle with metabolic dynamics and multiplexed imaging of iGlo with other biosensors of cell and metabolic state could unravel the causes of cellular heterogeneity in a single population. Amongst cell types, this variability in response likely reflects differences in how each cell type metabolizes glutamine. HeLa cells had significant changes in glutamine levels in both the cytoplasm and mitochondrial matrix when either glutamine synthesis or glutaminolysis were inhibited, whereas A549 cells appeared to have compartment-specific effects. In A549 cells, changes in glutamine dynamics in response to MSO and CB-839 were not significant in the mitochondrial matrix unlike the cytoplasm, which could indicate that the cytoplasmic pool of glutamine is more dynamic than the mitochondrial pool or be a consequence of the apparently lower dynamic range of iGlo in the mitochondrial matrix. Additionally, U2OS did not have significant changes at either location despite importing significant amounts of glutamine. This might reflect cellular need, where HeLa and A549 cells are likely using glutamine for processes like fueling the TCA cycle and ATP production, whereas U2OS cells could be using glutamine for processes not impacted by inhibition of glutamine synthetase or glutaminase, like protein synthesis^1^. U2OS cells are reported to direct glutamine towards production of asparagine using asparagine synthetase^42^. This metabolic preference of U2OS cells may in part explain our result, in which we observed no significant changes in glutamine following MSO treatment. Notably, our metabolomics results were overall supportive of our imaging results with iGlo. As live cell imaging with metabolite biosensors and metabolomics are complementary methods, the combination of these methods could be more widely employed to better understand spatiotemporal metabolic dynamics. Additionally, while not probed here, future work could involve a broader panel of pharmacological inhibitors targeting multiple glutamine-utilizing enzymes with iGlo to investigate cell type- and compartment-specific uses for glutamine beyond reactions catalyzed by glutamine synthetase and glutaminase.

Unlike HeLa, A549, and U2OS cells, the dynamics of HepG2 glutamine uptake and use were significantly suppressed when grown in DMEM, with large cell-to-cell heterogeneity. However, when grown in HPLM, we observed significant enhancement in glutamine uptake, as well as increased glutamine utilization at the single-cell level. Together, this data suggests that dynamics of metabolism, in particular glutamine metabolism, are significantly influenced by cell culture conditions. As many cell lines, particularly cancer cell lines, can adapt to survive under environmental changes, dynamic measurements using biosensors in single cells will reflect this^43^. Thus, our data suggests that careful consideration of culture conditions should be given when investigating and interpreting metabolic dynamics in single cells.

As glutamine and glucose metabolism are inextricably linked, we co-imaged iGlo with R-iLACCO1.2 to measure the dynamic interplay between glutamine use and lactate production in single HeLa cells. Using our multiplexing approach, we found that lactate production quickly increases when glutaminase is inhibited. This fast lactate accumulation seems to be due to a metabolic switch, as ATP levels remained consistent when measured using the ATP biosensor, iATPSnFR2^12^. To confirm our imaging data, we used respirometry and stable isotope tracing to find that in HeLa cells treated with CB-839, there is indeed a decrease in glutamine incorporation into the TCA cycle, decreased ATP production from oxidative phosphorylation, and an increase in glycolysis. Thus, we combined live, single cell imaging with readouts of cell function to uncover a rapid metabolic switch in HeLa cells. The combination of the two complimentary approaches allowed us to confirm a rapid switch from respiration to glycolysis to maintain energy homeostasis. Importantly, co-imaging of iGlo with R-iLACCO1.2 revealed the fast temporal dynamics of this metabolic switch, which might not otherwise be illuminated. By combining single cell measurements of glutamine dynamics with respirometry and metabolomics, we can measure both fast spatial and temporal changes in glutamine alongside larger-scale changes in metabolism, providing a holistic perspective into metabolism, which would not otherwise be possible.

Altogether, we have developed a new glutamine biosensor, iGlo, which we have used to measure the subcellular dynamics of glutamine metabolism. Using our biosensor, precise investigation into glutamine dynamics at subcellular level is possible, leading to a better understanding of metabolic regulation and adaptation in the cell.

## Methods

### Small molecules and plasmids

Ammonia chloride (NH_4_Cl, Fisher Scientific, Cat# A661-500) and methionine sulfoximine (MSO, Thermo Scientific, Cat# 227200010) were dissolved in PBS (Thermo Scientific Cat# 20012027). CB-839 (ThermoFisher Scientific, Cat# 50-163-7696) was dissolved in DMSO. L-glutamine (Thermo Scientific Cat# AAA1420130), L-alanine (Thermo Scientific Cat# AAA1580422), L-arginine (Thermo Fisher Scientific Cat# A15738-14), L-asparagine (Thermo Scientific Cat# AAB2147336), L-aspartic acid (Thermo Scientific Cat# AAA1352030), L-cysteine (Millipore Sigma Cat# 2430-M), L-glutamic acid (Thermo Scientific Cat #AC156212500), glycine (Sigma-Aldrich Cat# 50046), L-histidine (Sigma-Aldrich Cat# H-8125), L-isoleucine (Fisher Scientific Cat# BP384), L-leucine (Fisher Scientific Cat# BP385), L-lysine (Millipore Sigma Cat# 4400-M), L-methionine (Sigma-Aldrich Cat# M-9625), L-phenylalanine (Sigma-Aldrich Cat# P-2126), L-proline (Sigma-Alrich Cat# P-0380), L-serine (Sigma-Aldrich Cat# S4500), L-threonine (Sigma-Aldrich Cat# T-8625), L-tryptophan (Sigma-Aldrich Cat# T0254), L-tyrosine (Sigma-Aldrich Cat# T-3754), and L-valine (Fisher Scientific Cat# BP397) were dissolved in PBS.

pcDNA-R-iLACCO1.2 was a gift from Robert Campbell (Addgene plasmid # 208027 ; http://n2t.net/addgene:208027 ; RRID:Addgene_208027). pcDNA-R-diLACCO1 was a gift from Robert Campbell (Addgene plasmid # 208028 ; http://n2t.net/addgene:208028 ; RRID:Addgene_208028). pcDNA3.1(−) FLIPQTV3.0 2m (Addgene plasmid # 63731 ; http://n2t.net/addgene:63731 ; RRID:Addgene_63731) and pcDNA3.1(−) FLIPQTV3.0 100micro (Addgene plasmid # 63730 ; http://n2t.net/addgene:63730 ; RRID:Addgene_63730) were gifts from Sakiko Okumoto. pAAV-CAG-jAspSnFR3-mRuby3 was a gift from Lucas Sullivan (Addgene plasmid # 203459 ; http://n2t.net/addgene:203459 ; RRID:Addgene_203459). pAAV.CAG.(cyto).iATPSnFR2.A95K.HaloTag was a gift from Tim Brown & HHMI-JRC Tool Translation Team (Addgene plasmid # 209652 ; http://n2t.net/addgene:209652 ; RRID:Addgene_209652). pET28-MBP-super TEV protease was a gift from Mark Howarth (Addgene plasmid # 171782 ; http://n2t.net/addgene:171782 ; RRID:Addgene_171782). pcDNA 3.1(+) GlutaR was constructed through the Thermo Fisher Scientific GeneArt cloned genes service.

For cloning iGlo, GlnH was reconstructed from pcDNA3.1(−) FLIPQTV3.0 100micro by Gibson Assembly using GeneArt Gibson Assembly HiFi Master Mix (Thermo Fisher Scientific Cat# A46628)^44^. A Quikchange PCR was performed to reintroduce wild-type asparagine at GlnH position 157^45^. cpEGFP was amplified from Malibu, and superfolder mutations were introduced by Quikchange PCR^46,47^. The resulting cpsfEGFP variant, flanked by linkers, was then inserted into GlnH by Gibson Assembly. The annotated sequence of iGlo can be found in Supplementary Table 1. For making targeted iGlo, localization sequences were appended to the N-terminus of iGlo by Gibson Assembly. Targeting sequences can be found in Supplementary Table 2. diGlo was made using QuickChange PCR. To make mRuby3-iGlo, mRuby3 was amplified from pAAV-CAG-jAspSnFR3-mRuby3 and appended to the N-terminus of iGlo using Gibson Assembly.

### Screening biosensor variants in bacterial lysate

Plasmids were transformed into BL21 *E. coli* and plated on LB-ampicillin plates. iGlo variants were expressed in *E. coli* in either 1 mL (for linker screens) or 25 mL (for initial insertion site screen) cultures using autoinduction media^48^. Cultures were inoculated with a single colony and incubated with shaking at 220 rpm for 6 hours at 37°C followed by 20 hours at 20°C. Cultures were then pelleted by centrifugation. To reduce the residual endogenous glutamine levels, the pellets were resuspended in PBS for 1 hour with shaking at 20°C^49^. Cells were pelleted once again and resuspended in B-PER lysis buffer (Thermo Scientific Cat# P178248) containing protease inhibitors (Pierce Protease Inhibitor Tablets, Thermo Scientific Cat# A32963). The resulting lysate was clarified by centrifugation and the supernatant was used for subsequent fluorescence readings by plate reader (Molecular Devices SpectraMax iD5) in a 96-well assay plate (Genesee Scientific, Cat# 33−755). The plate reader was equipped with SoftMax Pro 7.1 data acquisition software (Molecular Devices). Fluorescence was measured with 405 nm and 480 nm excitation and 520 nm emission before and after adding glutamine to the lysate to a final concentration of 1 mM. ΔF/F_0_ was calculated by normalizing the difference between the final and initial fluorescence values to the initial fluorescence value.

### Linker screening

Linker screening was performed as previously described^46^. Briefly, site directed mutagenesis was performed using NNK degenerate codons to generate a Ser-X-X N-terminal linker library for the first round of linker screening, and an X-X C-terminal linker library for the second round of linker screening, where X can be any amino acid. Linker libraries were transformed into BL21 *E. coli* and colonies were grown on LB-ampicillin plates. For each round of linker screening, 192 fluorescent colonies were selected and lysate screening was performed as described in the previous section.

### Protein purification

BL21 *E. coli* containing plasmids for 6xHis-iGlo or 6xHis-diGlo (with TEV site) were grown in OM-I media and induced with 100 µM IPTG (Goldbio, Cat# I2481C50)^50^. After harvesting by centrifugation, cells were mechanically lysed using an EmulsiFlex-C3 (Avestin) in buffer (50 mM Tris, 150 mM NaCl, 20 mM imidazole, 5% glycerol, pH 8.0). The lysate was clarified by centrifugation. The supernatant was incubated with Ni-NTA resin (Thermo Scientific, Cat# 25215), washed, and eluted (50 mM Tris, 150 mM NaCl, 500 mM imidazole, 5% glycerol, pH 8.0). Eluate was concentrated using 10 kDa MWCO concentrators (Thermo Fisher Scientific Cat# 88528). MBP-super TEV protease (0.3 mg) was added to remove the His tag, and samples dialyzed with 10 kDa MWCO membrane (Fisher Scientific Cat# PI88243) into protein storage buffer (50 mM Tris, 150 mM NaCl, 5% glycerol, pH 8.0)^51^. Digested eluate was reapplied to Ni-NTA resin and washed. Flow-through and fractions were analyzed by SDS-PAGE (Thermo Fisher Scientific Cat# NP0322BOX) for purity. Final purified protein was frozen in liquid N□ and stored at −80□°C.

### In vitro iGlo assays

In vitro characterization of iGlo and diGlo was performed using opaque black 96-well plates (Genesee Scientific, Cat# 33−755). All plate reader-based in vitro assays utilized 6 µM protein in a protein assay buffer (50 mM Tris, 150 mM NaCl, 5% glycerol) at variable pH. For specificity and dose-response assays, an assay buffer with pH 7.2 was used. pH sensitivity assays were conducted with assay buffers adjusted to pH 6.5, 7.0, 7.2, 7.5, 7.7, and 8.0. K_D_ values were determined through nonlinear fitting using Equation 1 in GraphPad Prism 10.

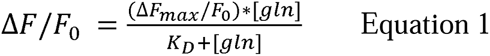

Fluorescence measurements for all assays were conducted from above, utilizing monochromators with excitation wavelengths of 405 nm and 480 nm and an emission wavelength of 520 nm. For excitation spectra, emission was detected at 560 nm. For emission spectra, excitation was conducted at 460 nm. Detection was performed using an ultra-cooled photomultiplier tube.

Stopped flow kinetics experiments were performed using an Applied Photometrics SX20. iGlo and glutamine in pH 8.0 protein assay buffer were mixed to a final concentration of 300 nM iGlo and variable glutamine concentrations ranging from 7.81 µM to 1000 µM. Experiments were conducted at 37 °C. An excitation wavelength of 480 nm was used and a bandpass of 6.98 nm was set using a monochromator (Applied Photophysics). Emission was filtered through a 515 nm longpass filter (CVI Laser). Fluorescence was measured every 360 ms for 360 seconds with detector voltage set to 300 V. Data was collected using the Pro-Data SX data acquisition software (Applied Photophysics). K_fast_ and K_slow_ values were determined through a nonlinear two-phase association fit (Equations 2a-2c) in GraphPad Prism 10.

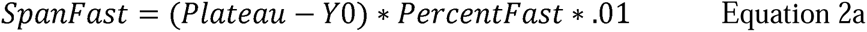

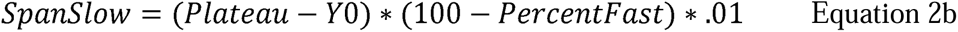

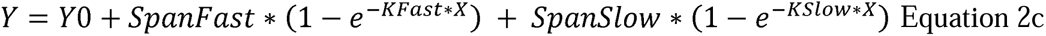

### Cell culture and transfection

HeLa (ATCC Cat# CRM-CCL-2), U2OS (ATCC Cat# HTB-96), A549 (ATCC Cat# CCL-185), and HepG2 (ATCC Cat# HB-8065) cells were cultured at 37□°C with 5% CO□ in DMEM (Thermo Scientific Cat# 10569010) containing 4.5□g/L glucose, 4 mM glutaMAX, sodium pyruvate, 10% FBS (Avantor 76419-584), and 100□U/mL penicillin-streptomycin (Fisher Scientific Cat# 15-140-122). Mycoplasma status was monitored by NucBlue staining (Fisher Scientific Cat# R37605) and PCR (Fisher Scientific Cat# AAJ66117AMJ).

HeLa and U2OS cells were plated at 150k–200k cells per dish one day before transfection or 75k–100k two days prior. A549 and HepG2 cells were plated at 75k–100k one day prior to transfection. Cells were seeded in 35 mm glass bottom dishes (Cellvis, Cat# d35-14-1.5-n). For experiments of HepG2 cells in HPLM, DMEM was exchanged for HPLM (Thermo Scientific Cat# A4899101), containing 5.0 mM glucose, 0.55 mM glutamine, 10% FBS, and 100 U/mL penicillin-streptomycin 24 hours after transfection, 24 hours prior to imaging. Transfections for HeLa and U2OS used 500 ng plasmid DNA, Opti-MEM (Gibco Cat #31985070), and FuGENE HD (Promega Cat# HD-1000) following manufacturer’s protocol and were conducted one day prior to imaging. Transfections for A549 were conducted similarly but with 1 µg plasmid DNA and performed two days prior to imaging. Transfections for HepG2 were conducted with 500□ng plasmid DNA and *Trans*IT-X2^®^ Dynamic Delivery System (Mirus, Cat# MIR 6003) two days prior to imaging.

### Fluorescence imaging and image analysis

Fluorescence microscopy experiments were conducted using a Nikon ECLIPSE Ti2 epifluorescence microscope with a CFI60 Plan Fluor 40x Oil Immersion Objective Lens (N.A. 1.3, W.D. 0.2 mm, F.O.V. 25 mm; Nikon), a Spectra III UV, V, B, C, T, Y, R, nIR light engine containing 380/20, 434/21, 475/28, and 575/25 LEDs, and Custom Spectra III filter sets (440/510/575 and 390/475/555/635/747) installed in a Ti cube polychroic. Imaging was captured with a Kinetix 22 back-illuminated sCMOS camera (Photometrics), and temperature was maintained at 37°C with a stage-top incubator (Tokai Hit). The system was operated with NIS-Elements software (Nikon). Exposure times ranged from 100 to 200 ms except for lyso-iGlo which 500 ms was used.

Cells were washed with HBSS (ThermoFisher Scientific Cat# 14065056 supplemented to 2 g/L glucose, 20 mM HEPES, pH 7.4) and incubated with HBSS for 25 minutes prior to imaging. For glutamine depleted conditions, 5 mM alanine was included in the HBSS during the 25-minute incubation and replaced with alanine-free HBSS immediately prior to imaging. Images were collected using a 40x objective. Images were taken every 30 seconds, and additions were made after 5 minutes of baseline imaging. For reversibility experiments, alanine was added at the 30 minute timepoint.

Image analysis was done as previously described using MATLAB R2024a (MathWorks)^46^. Cells or dishes that were not fully transfected, dying, out of focus, or under or over exposed were excluded from analysis.

### Confocal imaging

Confocal microscopy was used to image HeLa cells expressing iGlo localized to various subcellular compartments on a Leica Stellaris 5 DMi8 Confocal Microscope equipped with a DMOD WLL (440 nm – 790 nm), HC PL APO 63x/1.40 oil immersion CS2 lens, and a Power HyD S detector. LasX software (Leica) was used to control the microscope. Laser intensities between 0.5-8% were used to image cells. For cells expressing cytoplasmic and nuclear-localized iGlo, cells were washed with HBSS and stained with NucBlue (Fisher Scientific, Cat #R37605) for 5 minutes prior to imaging. Cells expressing mitochondrial matrix and OMM-iGlo were washed with HBSS and stained with 25 nM MitoTracker Deep Red (Thermo Scientific, Cat #M22426). Cells expressing iGlo targeted to the plasma membrane were washed with HBSS and stained with Deep Red CellMask Plasma Membrane Stain (Fisher Scientific, Cat#C10046). Cells expressing lysosomal iGlo were washed with HBSS and stained with 25 nM LysoTracker Deep Red (Thermo Scientific Cat #L12492). Cells were then treated with 1 mM glutamine (Sigma-Aldrich, Cat #607983) immediately prior to imaging.

### Flow cytometry

1.5×10^6^ HeLa cells were seeded in 10 cm cell culture dishes (Genesee Scientific Cat# 25-202). 24 hours later, transfection was performed with FuGENE HD (Promega Cat# HD-1000) following manufacturer’s protocol with 2.5 µg of mRuby3-iGlo or mRuby3-diGlo plasmid. 24 hours later, the cells were washed with HBSS (ThermoFisher Scientific Cat# 14065056 supplemented to 2 g/L glucose, 20 mM HEPES, pH 7.4) and incubated with 5 mM alanine in HBSS for 25 minutes. Cells were dissociated with TrypLE Express (ThermoFisher Scientific Cat# 12604013) and trypsin was inactivated with glutamine-free DMEM (ThermoFisher Scientific Cat# 12634010) containing 10% dialyzed FBS (Thermo Fisher Scientific Cat# 26-400-044), and 100□U/mL penicillin-streptomycin (Fisher Scientific Cat# 15-140-122). The cell suspension was centrifuged at 2000 g for 3 minutes and the supernatant was discarded. The cell pellet was resuspended in HBSS and separated into samples containing no glutamine or 1 mM glutamine.

The cell suspension was passed through a cell strainer (Fisher Scientific Cat# 08-771-23) immediately prior to loading onto a flow cytometer (Bio-Rad S3 Cell Sorter, Cat# 145-1002). The instrument was set to 37 °C and set to collect 50,000 events per run. Both lasers, 488 nm and 561 nm, were used. cpsfEGFP fluorescence was measured using 525/30 emission (FL1). mRuby3 fluorescence was measured using 586/25 emission (FL2). FlowJo v10 was used for analysis. A single population was observed and gated with FSC (forward scatter) and SSC (side scatter). The population was further gated using FL2, with gates carefully set such that there were a minimal number of events at the upper detection limit of FL1. Histograms showing gated FL1 fluorescence values were plotted in FlowJo v10 while the fluorescence values were exported and the means of FL1/FL2 ratio (cpsfEGFP/mRuby3 ratio) from biological replicates plotted in GraphPad Prism 10.

### LC-MS metabolomics

Mammalian cells were seeded into 6-well plates two days prior to drug treatment and each condition was harvested in triplicate. An additional well was cultured without treatment for packed cell volume (PCV) determination at the time of harvest. On day of harvest, the additional well was briefly washed with HBSS (ThermoFisher Scientific Cat# 14065056 supplemented to 2 g/L glucose, 20 mM HEPES, pH 7.4), incubated with trypsin, neutralized with DMEM, and pipetted into PCV tubes for centrifugation at 2.5k rcf for 1 min. The treatment of cells for metabolomics was performed identically to imaging conditions, with a 25 min preincubation in HBSS at 37 °C. Cells for NH_4_Cl and CB-839 experiments were pretreated with 5 mM alanine during the preincubation. Cells were treated with 200 μM NH_4_Cl or PBS control for 25 minutes, 5 mM methionine sulfoximine (MSO) or PBS control for 55 minutes, or 10 μM CB-839 or DMSO control for 115 minutes. Cells were incubated in HBSS at 37 °C during treatment. To harvest cells, a ratio of 50:1 of 80% MeOH to packed cell volume was used. Cells were incubated with 80% MeOH for 10 minutes on dry ice before collected using cell scrapers (Genesee Scientific Cat# 25-270) and transferred to tubes on dry ice. Cell extract was vortexed for 20 seconds prior to centrifugation at 16,000 rcf for 10 minutes at 4 °C. A second centrifugation step of the supernatant was performed under identical conditions for 10 minutes to remove residual cellular debris. The clarified supernatant was transferred to LC-MS vials and stored at −80°C until LC-MS analysis. Quality control samples were prepared by pooling equal volumes from each replicate.

Metabolites were separated using a Waters BEH z-HILIC column (150 × 2.1 mm, 2.7 µm) connected to a Thermo Scientific™ Orbitrap Exploris™ 480 mass spectrometer with a heated electrospray ionization (H-ESI) source. In positive ionization mode, the spray voltage was set to 3.5 kV, the heated capillary temperature to 320°C, the sheath gas flow to 35, the auxiliary gas flow to 10, and the sweep gas flow to 1. The same gas-flow parameters were used in negative-ionization mode, but with a spray voltage of 2.5 kV. The column temperature was maintained at 25°C, and the autosampler at 4°C. The mobile phase consisted of Solvent A: 10 mM ammonium bicarbonate in water (pH ∼9.1) and Solvent B: 95% acetonitrile/5% water. Chromatographic separation was conducted at 0.18 mL/min with the following gradient: 0-1 min, 95% B; 1-15 min, linear decrease to 45% B; 15-19 min, 45% B; 19-22 min, ramp back to 95% B; 22-34.5 min, 95% B for re-equilibration.

MS data was acquired using data-dependent acquisition (DDA) mode with polarity switching enabled. Full MS1 scans were collected at 120,000 resolution (m/z 70-1000) with an AGC target of 1 × 10^6. Raw LC-MS data was converted to mzXML format with MSConvert and processed in El MAVEN (v0.12.0). Peak detection, chromatogram alignment, and automated isotope annotation were performed using default settings. Retention times were normalized with external standards, ion intensities were corrected for matrix effects, and peaks were manually curated as needed. Processed peak tables were exported in CSV format for analysis and relative metabolite quantification.

### Seahorse XF analysis

All oxygen consumption and extracellular acidification measurements were performed on XF96 Agilent Analyzer. HeLa cells were plated at 2×10^4^ cells/well 24 hours prior to the experiment. Assays were performed at 37°C in DMEM medium (DMEM Sigma #5030, 31.6 mM NaCl, 3mg/L phenol red, and 5 mM HEPES) supplemented with 8 mM glucose, 2 mM glutamine, and 2 mM pyruvate at pH 7.4. 10 μM CB839 was added acutely 15 minutes prior to the initial measurements. Respiratory parameters were determined based on measurements taken in response to the following compounds: 2.25 μM oligomycin, 750 nM FCCP, 200 nM rotenone, 1 μM antimycin A. Respiratory parameters and rates of total ATP production were calculated as previously described^52,53^.

### Stable isotope tracing

HeLa cells were plated at 5×10^4^ cells/well in a 12-well cell culture plate in DMEM (Gibco 11965) supplemented with 2 mM GlutaMAX, 1% (v/v) penicillin/streptomycin, and 10% (v/v) FBS (all supplements from Gibco). After 24 hours, cell culture medium was replaced with medium for stable isotope tracing containing either 10 mM ^13^C_6_-glucose (CLM-1396) or 2 mM ^13^C_5_-glutamine (CLM-1822), as well as +/− 2 μM CB839. 24 hours after stable isotope tracing +/−CB839, cells were washed with ice cold 0.9% (w/v) saline prior to a Folch-like extraction using 5:2:5 ratio of methanol:water:chloroform. The water contained 1 μg of norvaline used as an internal standard. Samples were then quickly vortexed and centrifuged at 10,000*g* for 6 min at 4°C. The top, polar layer was collected and dried overnight in a refrigerated centrivap. Polar metabolites were derivatized using 20 μL of 2% (w/v) methoxyamine in pyridine for 45 min at 37°C, followed by incubation with 20 μL of MTBSTFA + 1% TBDMS (N-tert-Butyldimethylsilyl-N-methyltrifluoroacetamide with 1% tertButyldimethylchlorosilane) for 45 min at 37°C. Samples were analyzed via GC/MS using Agilent MassHunter software and Agilent Technologies DB-35 column as previously described^54^.

### Statistics and reproducibility

Figure preparation and statistical analysis were performed using GraphPad Prism 10, except for Fig S1c-d which were prepared in FlowJo v10. The statistical tests used and P values are reported in all figure legends where applicable. The number of cells analyzed (n cells), and number of independent experiments are reported in all figure legends. For imaging experiments, statistics were performed using the means of all single cell measurements from independent experiments. All time courses shown are the mean plus all cells unless otherwise noted. All dot plots shown depict the mean□±□standard deviation. When applicable, all data was plotted using the SuperPlots method^55^.

## Code and Data Availability Statement

Custom MATLAB code used to calculate t_1/2_ has been uploaded to GitHub (https://github.com/dlschmitt/t1-2_calculation-for-Scully-et-al) along with a readme file and sample data. Plasmids for iGlo have been deposited with Addgene. Metabolomics data will be deposited with Metabolomics Workbench.

## Author Contributions

J.M.S. and D.L.S. conceived the project. J.M.S and T.D.R. developed iGlo. J.M.S., M.J.S., and S.D. performed and analyzed all time course imaging. N.J.A. performed respirometry and stable isotope tracing. J.M.S., T.S.D, and E.D.K performed and analyzed all other metabolomic experiments. K.G.V. performed confocal microscopy. D.L.S., A.S.D., and T.T. oversaw all experiments. J.M.S., N.J.A., and D.L.S. prepared the manuscript and figures. All authors contributed to and approved the final version of the manuscript.

## Acknowledgements

We thank Peter DePaola IV for his help with protein purification, Mark Arbing for his assistance with flow cytometry, Martin Phillips for his assistance with stopped flow, and Katarina A. Cohen for her assistance with cell culture. We thank Michael Lawson for material support. We thank the UCLA-DOE Protein Expression Technology Center for access to a Bio-Rad S3 cell sorter, which is supported by Department of Energy Grant DE-FC02-02ER63421. We thank the UCLA Biochemistry Instrumentation Facility for access to a stopped flow apparatus, which was supported by the NIH (1S10OD023584). This work was supported by the National Institutes of Health (NIH/NIGMS T32GM145388 to J.M.S. and K.G.V., P30CA016042, R35GM138003 to A.D.S, and DP2GM154012-01 to D.L.S.); the UCLA Jonsson Comprehensive Cancer Center Strategic Plan Aligned Grant (to D.L.S.) and Graduate Research Award (to J.M.S. and T.S.D.): the UCLA Society of Hellman Fellows (to D.L.S.); and the Chan-Zuckerberg Initiative (MET-000000000151 to T.T. and D.L.S.).

