## Supplemental Information for "Illuminating spatial dynamics of glutamine metabolism with a sensitive genetically encoded biosensor"

**Supplemental Figures and Tables**

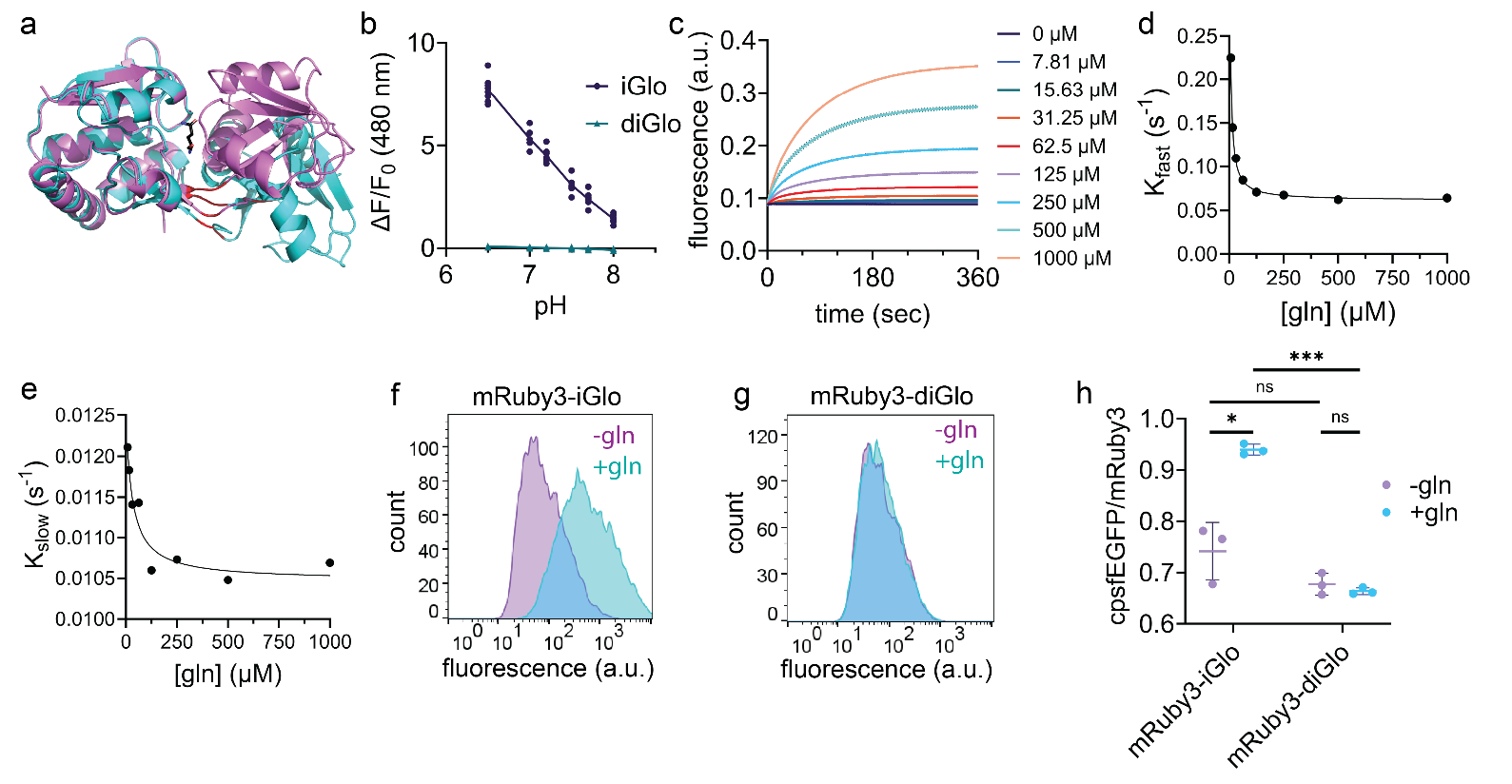

**Figure S1. Development and characterization of iGlo**

a, Aligned structures of glutamine-bound GlnH (violet, PDB 1WDN) and glutamine-free GlnH (cyan, PDB 1GGG). Sites where cpsfEGFP was inserted are shown in red. Glutamine is shown in black.

b, Effect of pH on the fractional change in fluorescence (ΔF/F_0_) of iGlo and diGlo in response to 1 mM glutamine in vitro. n = 8 from two independent protein preparations.

c, Fluorescence vs. time of 300 nM iGlo mixed with the indicated concentrations of glutamine measured by stopped flow. Solid lines indicate the means of 3-4 technical replicates and dots above and below the lines indicate mean ± S.D. Traces were fit using two phase association in GraphPad Prism 10 to obtain K_fast_ and K_slow_ values.

d-e, K_fast_ (d) and K_slow_ (e) vs. glutamine concentration of the fluorescence increase following mixing of 300 nM iGlo with glutamine measured by stopped flow. Curves shown are from a three parameter fit to the Hill equation performed in GraphPad Prism 10.

f-g, Representative flow cytometry measurements of HeLa cells in suspension expressing mRuby3-iGlo (c) and mRuby3-diGlo (d) with no glutamine (purple) or 1 mM glutamine (teal) showing the distribution of cpsfEGFP fluorescence of gated cells. Cells were gated using mRuby3 fluorescence as an expression marker. Adherent cells were incubated with 5 mM alanine for 25 minutes prior to dissociating with trypsin. FL1 corresponds to fluorescence measured using a 525/30 nm emission filter.

h, Mean cpsfEGFP/mRuby3 ratios of mRuby3-iGlo and mRuby3-diGlo HeLa cells measured by flow cytometry with no glutamine (purple) or 1 mM glutamine (cyan) across three biological replicates. mRuby3-iGlo * P = 0.019; mRuby3-diGlo n.s. P = 0.29; -gln n.s. P = 0.10; +gln *** P = 0.0001 by two-tailed paired t test. Mean ± S.D. is shown.

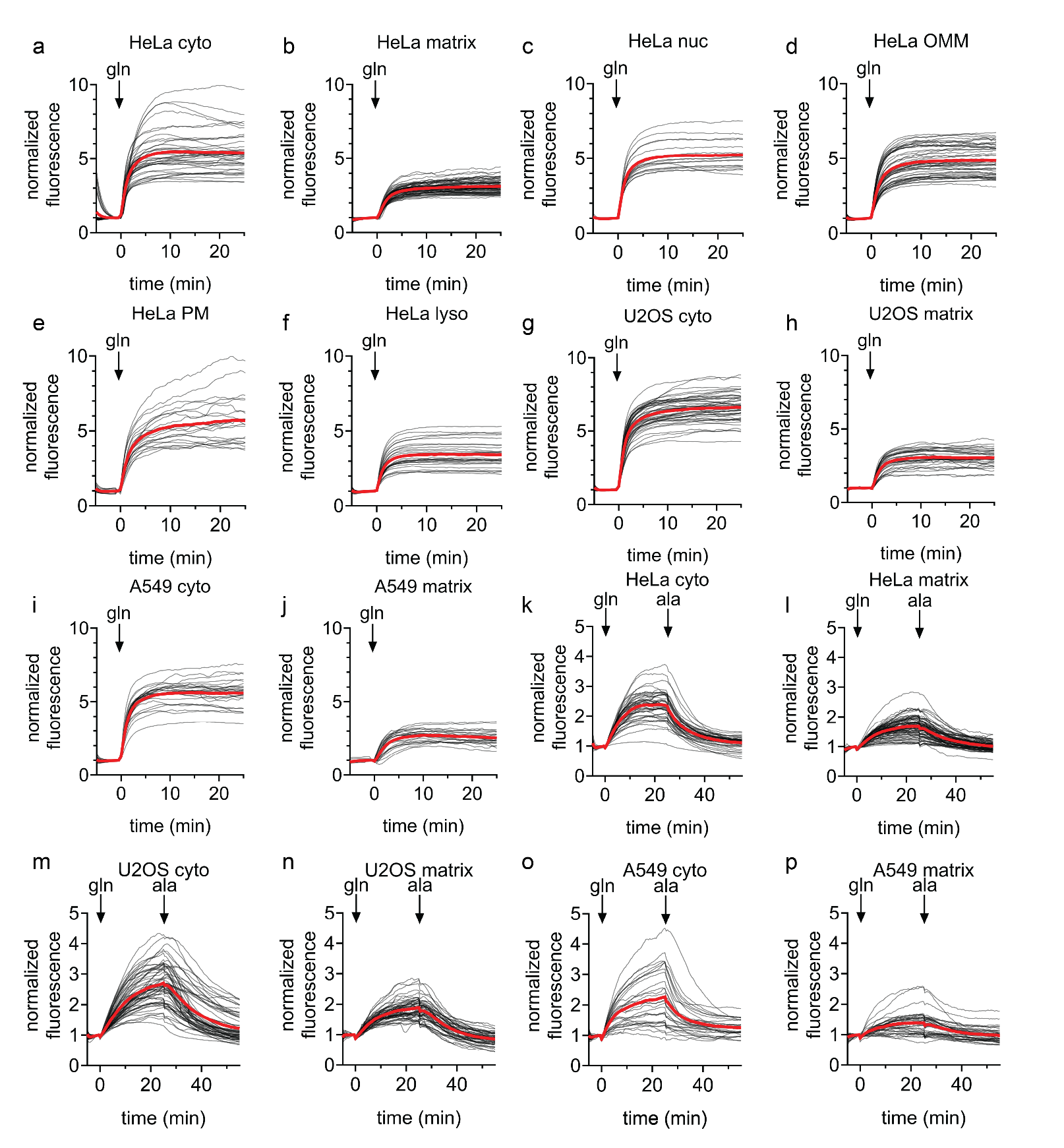

**Figure S2. Measurement of subcellular glutamine import and export with iGlo**

a-f, Mean (red) and single cell (black) normalized fluorescence of cyto-iGlo (a, n = 39 cells), matrix-iGlo (b, n = 71 cells), nuc-iGlo (c, n = 16 cells), OMM-iGlo (d, n = 48 cells), PM-iGlo (e, n = 21 cells), and lyso-iGlo (f, n = 28 cells) expressed in HeLa cells following the addition of 1 mM glutamine across three biological replicates. Cells were treated with 5 mM alanine prior to imaging for 25 minutes.

g-h, Mean (red) and single cell (black) normalized fluorescence of cyto-iGlo (g, n = 46 cells) and matrix-iGlo (h, n = 33 cells) expressed in U2OS cells following the addition of 1 mM glutamine three biological replicates. Cells were treated with 5 mM alanine prior to imaging for 25.

i-j, Mean (red) and single cell (black) normalized fluorescence of cyto-iGlo (I, n = 25 cells) and matrix-iGlo (j, n = 21 cells) expressed in A549 cells following the addition of 1 mM glutamine three biological replicates. Cells were treated with 5 mM alanine prior to imaging for 25.

k-l, Mean (red) and single cell (black) normalized fluorescence of cyto-iGlo (k, n = 43 cells) and matrix-iGlo (l, n = 77 cells) expressed in HeLa cells following the addition of 1 µM glutamine and 5 mM alanine across three biological replicates. Cells were treated with 5 mM alanine prior to imaging for 25.

m-n, Mean (red) and single cell (black) normalized fluorescence of cyto-iGlo (m, n = 58 cells) and matrix-iGlo (n, n = 54 cells) expressed in U2OS cells following the addition of 1 µM glutamine and 5 mM alanine across three biological replicates. Cells were treated with 5 mM alanine prior to imaging for 25.

o-p, Mean (red) and single cell (black) normalized fluorescence of cyto-iGlo (o, n = 28 cells) and matrix-iGlo (p, n = 36 cells) expressed in A549 cells following the addition of 1 µM glutamine and 5 mM alanine across three biological replicates. Cells were treated with 5 mM alanine prior to imaging for 25.

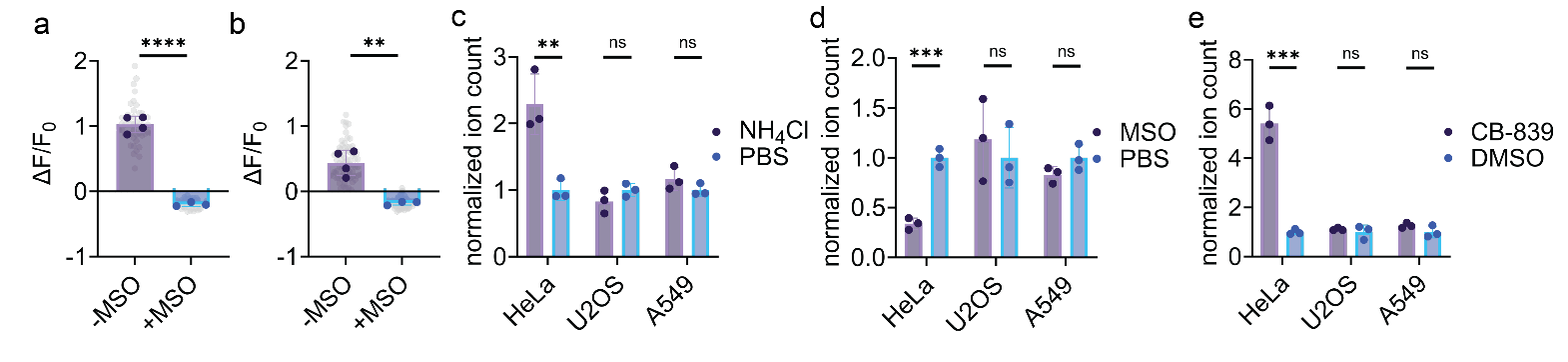

**Figure S3. iGlo reveals real-time dynamics of glutamine metabolism**

a-b, Fractional change in fluorescence (ΔF/F_0_) of HeLa cells expressing cyto-iGlo (a) or matrix-iGlo (b) after 25 minutes following the addition of 200 µM NH_4_Cl in the absence (purple; cyto-iGlo n = 46 cells from 4 independent experiments; matrix-iGlo n = 77 cells from 4 independent experiments) or presence (blue; cyto-iGlo n = 77 cells from 3 independent experiments; matrix-iGlo n = 97 cells from 3 independent experiments) of 5 mM methionine sulfoximine (MSO). Cells were treated with 5 mM alanine prior to imaging for 25 minutes to deplete intracellular glutamine. Grey dots represent single cell measurements. Solid dots represent the mean of single cell measurements from biological replicates. Cyto (a) **** P < 0.0001; matrix (b) ** P = 0.0029 by unpaired t test.

c, Quantification of glutamine levels by LC-MS in HeLa, U2OS, and A549 cells following treatment with 200 µM NH_4_Cl relative to vehicle (PBS). Cells were treated with 5 mM alanine for 25 minutes prior to NH_4_Cl or PBS addition. HeLa ** P = 0.0096; U2OS n.s. P = 0.21; A549 n.s. P = 0.22 by unpaired two-tailed t test.

d, Quantification of glutamine levels by LC-MS in HeLa, U2OS, and A549 cells following treatment with 5 mM methionine sulfoximine (MSO) relative to vehicle (PBS). HeLa *** P = 0.0004; U2OS ** P = 0.0088; A549 n.s. P = 0.14 by unpaired two-tailed t test.

e, Quantification of glutamine levels by LC-MS in HeLa, U2OS, and A549 cells following treatment with 10 µM CB-839 relative to vehicle (DMSO). Cells were treated with 5 mM alanine for 25 minutes prior to CB-839 or DMSO addition. HeLa *** P = 0.0004; U2OS n.s. P = 0.55; A549 n.s. P = 0.16 by unpaired two-tailed t test.

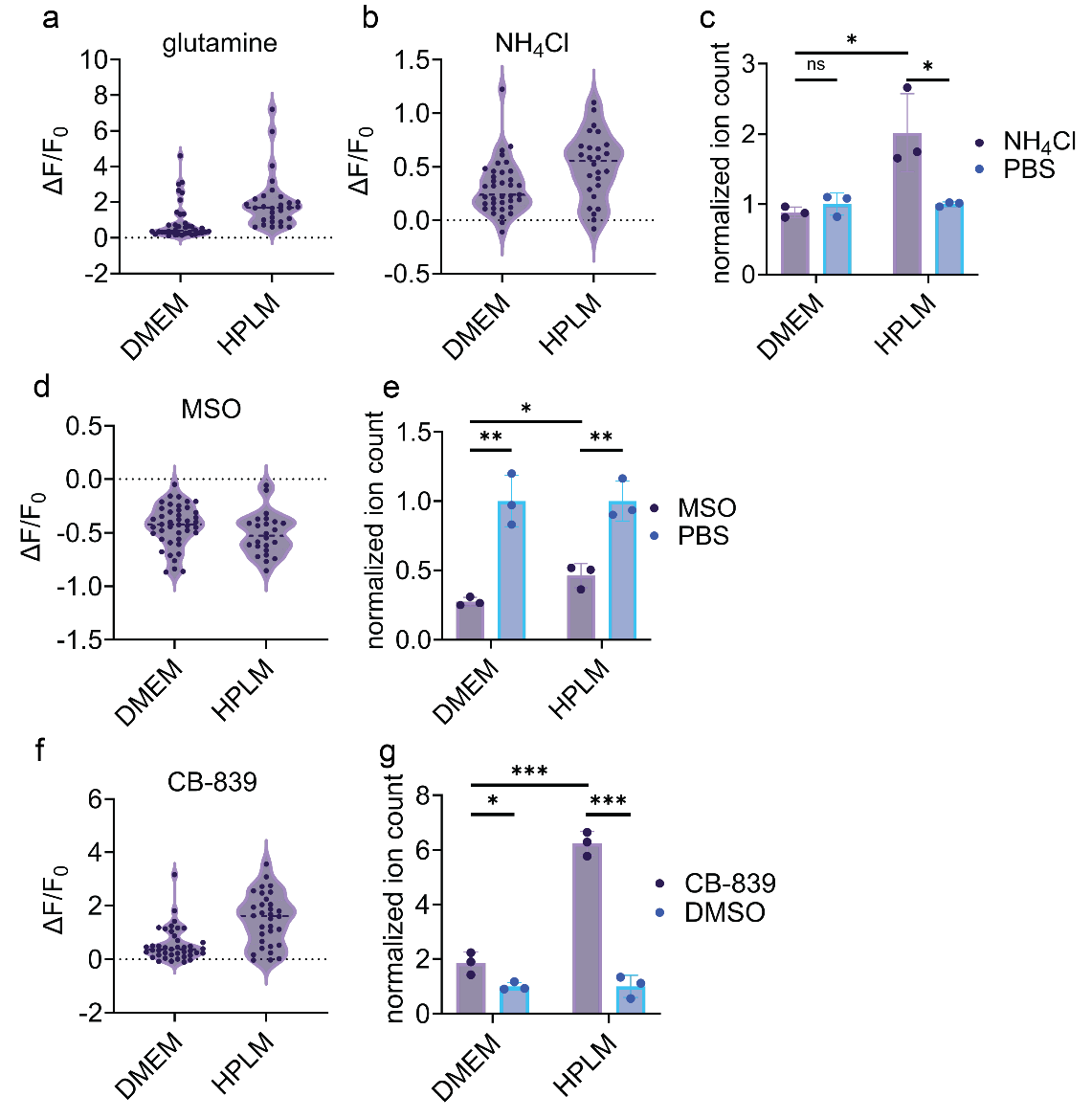

**Figure S4. Culture conditions impact glutamine metabolism in HepG2 cells**

a, Distribution of fractional changes of fluorescence (ΔF/F_0_) of single HepG2 cells either cultured in DMEM only or HPLM 24 hours prior to imaging and treated with 1 mM glutamine**.**

b, Distribution of fractional changes of fluorescence (ΔF/F_0_) of single HepG2 cells either cultured in DMEM only or HPLM 24 hours prior to imaging treated with 200 µM NH_4_Cl**.**

c, Quantification of glutamine levels by LC-MS in HepG2 cells either cultured in DMEM only or HPLM 24 hours following treatment with 200 µM NH_4_Cl relative to vehicle (PBS). Cells were treated with 5 mM alanine for 25 minutes prior to NH_4_Cl or PBS addition. DMEM n.s. P = 0.31; HPLM ** P = 0.034; DMEM v HPLM * P = 0.025 by unpaired two-tailed t test.

d, Distribution of fractional changes of fluorescence (ΔF/F_0_) of single HepG2 cells either cultured in DMEM only or HPLM 24 hours prior to imaging treated with 5 mM MSO**.**

e, Quantification of glutamine levels by LC-MS in HepG2 cells either cultured in DMEM only or HPLM 24 hours following treatment with 5 mM MSO relative to vehicle (PBS). DMEM ** P = 0.0026; HPLM ** P = 0.0050; DMEM v HPLM * P = 0.024 by unpaired two-tailed t test.

f, Distribution of fractional changes of fluorescence (ΔF/F_0_) of single HepG2 cells either cultured in DMEM only or HPLM 24 hours prior to imaging treated with 10 µM CB-839**.**

g, Quantification of glutamine levels by LC-MS in HepG2 cells either cultured in DMEM only or exchanged for HPLM 24 hours following treatment with 10 µM CB-839 relative to vehicle (DMSO). Cells were treated with 5 mM alanine for 25 minutes prior to CB-839 or DMSO addition. DMEM * P = 0.027; HPLM *** P = 0.0001; DMEM v HPLM *** P = 0.0002 by unpaired two-tailed t test.

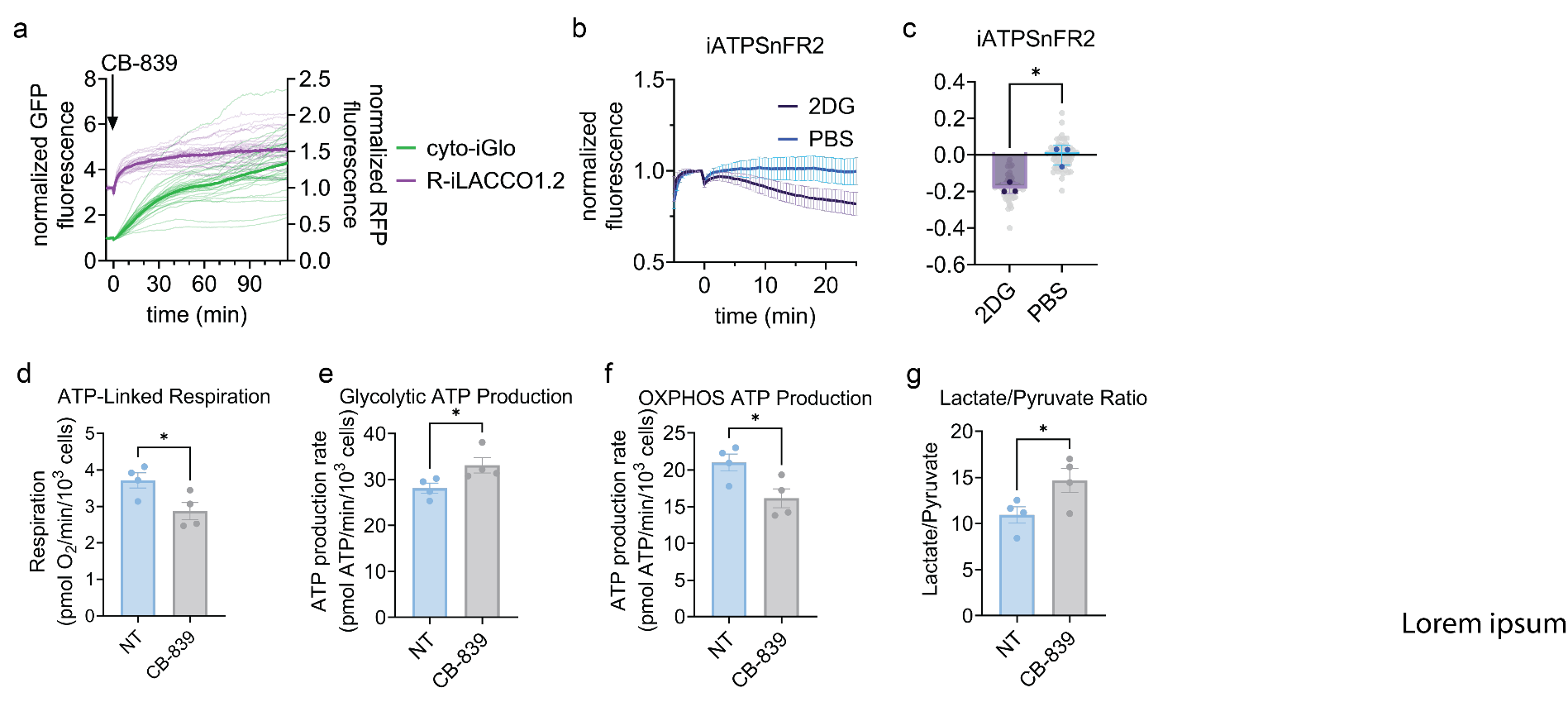

**Figure S5. Multiplexed imaging with iGlo reveals rapid metabolic adaptations**

a, Normalized fluorescence of single cells (thin lines) and mean normalized fluorescence (thick lines) of HeLa cells expressing cyto-iGlo (green) and R-iLACCO1.2 (magenta) following addition of 10 µM CB-839 across three biological replicates. Cells were treated with 5 mM alanine prior to imaging for 25 minutes. n = 30 cells.

b, Mean normalized fluorescence of HeLa cells expressing iATPSnFR2 treated with 40 mM 2-deoxy-D-glucose (2DG, purple, n = 60 cells) or PBS (blue, n = 60 cells) across three biological replicates. Mean ± S.D. is shown.

c, ΔF/F_0_ of HeLa cells expressing iATPSnFR2 after 25-minute treatments with 2-deoxy-D-glucose (2DG, purple, n = 60 cells) or PBS (blue, n = 60 cells). Grey dots represent single cell measurements and solid dots represent the means of the three biological replicates. * P = 0.021 by unpaired two-tailed t test.

d, ATP-linked respiration of HeLa cells following treatment with 10 µM CB-839 or no treatment. * P = 0.014 by paired two-tailed t test.

e, ATP production from glycolysis in HeLa cells following treatment with 10 µM CB-839 or no treatment. * P = 0.028 by paired two-tailed t test.

f, ATP production from OXPHOS in HeLa cells following treatment with 10 µM CB-839 or no treatment. * P = 0.0109 by paired two-tailed t test.

g, Lactate/pyruvate ratio in HeLa cells following 24-hour treatment with 2 µM CB-839 or no treatment. * P = 0.0146 by paired two-tailed t test.

**Table S1. Protein sequences**

| Protein Name | Amino acid sequence  Blue; *E. coli* GlnH  Orange; Arg-75 of GlnH (Met in diGlo)  Red; linkers  Green; cpsfEGFP |
| --- | --- |
| iGlo | MAGSKLVVATDTAFVPFEFKQGDLYVGFDVDLWAAIAKELKLDYELKPMDFSGIIPALQTKNVDLALAGITITDERKKAIDFSDGYYKSSVCNVYIMADKQKNGIKANFKIRHNVEDGSVQLADHYQQNTPIGDGPVLLPDNHYLSTQSVLSKDPNEKRDHMVLLEFVTAAGITLGMDELYKGGTGGSMSKGEELFTGVVPILVELDGDVNGHKFSVRGEGEGDATNGKLTLKFICTTGKLPVPWPTLVTTLTYGVQCFSRYPDHMKQHDFFKSAMPEGYVQERTISFKDDGTYKTRAEVKFEGDTLVNRIELKGIDFKEDGNILGHKLEYNNSGLLVMVKANNNDVKSVKDLDGKVVAVKSGTGSVDYAKANIKTKDLRQFPNIDNAYMELGTNRADAVLHDTPNILYFIKTAGNGQFKAVGDSLEAQQYGIAFPKGSDELRDKVNGALKTLRENGTYNEIYKKWFGTEPK |
| diGlo | MAGSKLVVATDTAFVPFEFKQGDLYVGFDVDLWAAIAKELKLDYELKPMDFSGIIPALQTKNVDLALAGITITDEMKKAIDFSDGYYKSSVCNVYIMADKQKNGIKANFKIRHNVEDGSVQLADHYQQNTPIGDGPVLLPDNHYLSTQSVLSKDPNEKRDHMVLLEFVTAAGITLGMDELYKGGTGGSMSKGEELFTGVVPILVELDGDVNGHKFSVRGEGEGDATNGKLTLKFICTTGKLPVPWPTLVTTLTYGVQCFSRYPDHMKQHDFFKSAMPEGYVQERTISFKDDGTYKTRAEVKFEGDTLVNRIELKGIDFKEDGNILGHKLEYNNSGLLVMVKANNNDVKSVKDLDGKVVAVKSGTGSVDYAKANIKTKDLRQFPNIDNAYMELGTNRADAVLHDTPNILYFIKTAGNGQFKAVGDSLEAQQYGIAFPKGSDELRDKVNGALKTLRENGTYNEIYKKWFGTEPK |

**Table S2. Localization sequences**

| cyto-iGlo  N-terminal localization sequence | MLPPLERLTL |
| --- | --- |
| matrix-iGlo  N-terminal localization sequence | MSVLTPLLLRGLTGSARRLPVPRAKIHSLGDPMSVLTPLLLRGLTGSARRLPVPRAKIHSLGDP |
| nuc-iGlo  N-terminal localization sequence | MPKKKRKVPKKKRKV |
| OMM-iGlo  N-terminal localization sequence | MAIQLRSLFPLALPGMLALLGWWWFFSRKK |
| lyso-iGlo  N-terminal localization sequence | MAAPGSARRPLLLLLLLLLLGLMHCASAAMFMVKNGNGTACIMANFSAAFSVNYDTKSGPKNMTFDLPSDATVVLNRSSCGKENTSDPSLVIAFGRGHTLTLNFTRNATRYSVQLMSFAYNLSDTHLFPNASSKEIKTVESITDIRADIDKKYRCVSGTQVHMNNVTVTPHDATIQAYLSNSSFSRGETRCEQDRPSPTTAPPAPPSPSPSPVPKSPSVDKYNVSGTNGTCLLASMGLQLNLTYERKDNTTVTRLLNINPNKTSASGSCGAHLVTLELHSEGTTVLLFQFGMNASSSRFFLQGIQLNTILPDARDPAFKAANGSLRALQATVGNSYKCNAEEHVRVTKAFSVNIFKVWVQAFKVEGGQFGSVEECLLDENSMLIPIAVGGALAGLVLIVLIAYLVGRKRSHAGYQTI |
| PM-iGlo  N-terminal localization sequence | MGCIKSKRKDKGG |
